# Structure-specific variation in per- and polyfluoroalkyl substances toxicity among genetically diverse *Caenorhabditis elegans* strains

**DOI:** 10.1101/2024.05.29.596269

**Authors:** Tess C. Leuthner, Sharon Zhang, Brendan F Kohrn, Heather M. Stapleton, L. Ryan Baugh

## Abstract

**Background:** There are >14,500 structurally diverse per- and polyfluoroalkyl substances (PFAS). Despite knowledge that these “forever chemicals” are in 99% of humans, mechanisms of toxicity and adverse health effects are incompletely known. Furthermore, the contribution of genetic variation to PFAS susceptibility and health consequences is unknown.

**Objectives:** We determined the toxicity of a structurally distinct set of PFAS in twelve genetically diverse strains of the genetic model system *Caenorhabditis elegans*.

**Methods:** Dose-response curves for four perfluoroalkyl carboxylic acids (PFNA, PFOA, PFPeA, and PFBA), two perfluoroalkyl sulfonic acids (PFOS and PFBS), two perfluoroalkyl sulfonamides (PFOSA and PFBSA), two fluoroether carboxylic acids (GenX and PFMOAA), one fluoroether sulfonic acid (PFEESA), and two fluorotelomers (6:2 FCA and 6:2 FTS) were determined in the *C. elegans* laboratory reference strain, N2, and eleven genetically diverse wild strains. Body length was quantified by image analysis at each dose after 48 hr of developmental exposure of L1 arrest-synchronized larvae to estimate effective concentration values (EC50).

**Results:** There was a significant range in toxicity among PFAS: PFOSA > PFBSA ≈ PFOS ≈ PFNA > PFOA > GenX ≈ PFEESA > PFBS ≈ PFPeA ≈ PFBA. Long-chain PFAS had greater toxicity than short-chain, and fluorosulfonamides were more toxic than carboxylic and sulfonic acids. Genetic variation explained variation in susceptibility to PFBSA, PFOS, PFBA, PFOA, GenX, PFEESA, PFPeA, and PFBA. There was significant variation in toxicity among *C. elegans* strains due to chain length, functional group, and between legacy and emerging PFAS.

**Conclusion:** *C. elegans* respond to legacy and emerging PFAS of diverse structures, and this depends on specific structures and genetic variation. Harnessing the natural genetic diversity of *C. elegans* and the structural complexity of PFAS is a powerful New Approach Methodology (NAM) to investigate structure-activity relationships and mechanisms of toxicity which may inform regulation of other PFAS to improve human and environmental health.

## Introduction

Per- and polyfluoroalkyl substances (PFAS) have been used widely in commercial products and industrial processes since the mid-1900s (Lindstrom et al. 2011). Given their surfactant properties, these compounds have largely been used in stain and water-resistant materials such as non-stick cookware, food packaging (*e.g.* Teflon® from 3M), and fabrics (*e.g.* waterproof clothes and furniture), as well as industrial applications including hydraulic fluids and aqueous fire-fighting foams, and many other products (Glüge et al. 2020). The abundant use of PFAS and their commercial success can be attributed to their physiochemical properties resulting from strong carbon-fluorine bonds. However, these properties make many PFAS highly bioaccumulative, mobile and persistent, with no estimated half-lives. This has resulted in ubiquitous contamination of water, air, and soil (Evich et al. 2022). These “forever chemicals” are detected globally in drinking water and food sources (plants and animals), which are both major routes of human exposure (Ghisi et al. 2019; Salvatore et al. 2022). In fact, 99% of all human serum samples tested in the United States contain PFAS (Graber et al. 2019; Kato et al. 2011).

Only a few of the >14,500 structurally diverse PFAS have been tested for safety (United States Environmental Protection Agency 2021). Two well-studied PFAS, perfluorooctanoic acid (PFOA) and perfluorooctanesulfonic acid (PFOS) are now phased out of intentional production and considered “legacy” PFAS chemicals. These long-chain PFAS (≥ 7 and ≥ 6 perfluorinated carbons for carboxylic and sulfonic acids, respectively) have been replaced with short-chain compounds (6 or less and 5 or less perfluorinated carbons for carboxylic and sulfonic acids, repsectively), or substances that have ether linkages in the carbon backbone (such as GenX) (Buck et al. 2011). These “emerging” PFAS, particularly short-chain and ultrashort PFAS chemicals, now make up a majority of the PFAS in consumer and industrial products, the environment, and people (McCord and Strynar 2019; Zheng et al. 2023). However, the mechanisms of toxicity and impacts on health of most PFAS remain unknown (Pelch et al. 2022). With so many PFAS chemicals, it is imperative to systematically evaluate structure-activity relationships, but only a few studies have investigated how molecular structures drive PFAS toxicity, and very few studies have compared toxicity between legacy and emerging PFAS (Conley et al. 2022, 2024; Cope et al. 2021; Gebreab et al. 2020; Truong et al. 2022). This is a major public and environmental health crisis, as PFAS exposures are associated with increased cancer incidence (such as breast, kidney, prostate, and testicular cancers) in addition to adverse effects on lipid metabolism and reproductive, developmental, hepatic, thyroid, renal, and immune systems (Birru et al. 2021; Bonefeld-Jorgensen et al. 2011; Fenton et al. 2021; Hall et al. 2022, 2023; Mancini et al. 2020; Perng et al. 2022; Radke et al. 2022; Shearer et al. 2021; Tsai et al. 2020; Wielsøe et al. 2017).

Human populations are diverse, but a missing, yet critical aspect in risk assessment is how genetic variation contributes to variation in susceptibility and response to exposures (Zeise et al. 2013). The use of mutant mouse strains for structure-activity relationship studies was instrumental in regulating another complex class of pollutants called dioxins and dioxin-like compounds (Poland and Glover 1974; Birnbaum et al. 1990). The invaluable Mouse Diversity Panel Resources such as the Collaborative Cross and Diversity Outbred models have also demonstrated the contribution of genetic variation to toxicity, mainly of a few known carcinogens such as trichloroethylene, benzene, and arsenic (French et al. 2015; Harrill and McAllister 2017; Stýblo et al. 2019; Venkatratnam et al. 2017). These and other foundational studies demonstrate the power of harnessing genetic variation in model organisms to analyze toxicant structure-activity relationships and set a precedent for informing regulatory policy (Bălan et al. 2021; Cousins et al. 2020; Kwiatkowski et al. 2020). Similar work is needed to determine if it is appropriate to regulate the thousands of PFAS chemicals in production today as a single chemical class.

Due to its fast growth rate, short generation time, large broods, ease of liquid culture, and developmental synchronization in the first larval stage (L1 arrest), *C. elegans* is highly amenable to toxicity assays (Boyd et al. 2010; Hartman et al. 2021; Hibshman et al. 2021; Hunt 2016; Leung et al. 2008). Furthermore, the National Institutes of Health goal of refining, reducing, and replacing vertebrate animal testing in toxicology, along with the impracticality of testing >14,500 compounds, makes *C. elegans* an ideal model organism for investigating effects of PFAS and other pollutants on health and disease (Collins et al. 2008; Grimm 2019; Krewski et al. 2020; Stokes 2015). Importantly, *C. elegans* has a similar number of protein-coding genes as humans, ∼60-80% of which are homologous, and it has comparable developmental responses to ToxCast chemicals as zebrafish, rats, and rabbits (Boyd et al. 2016). *C. elegans* has been instrumental in numerous fundamental advances in genetics, developmental biology, and neurobiology. Groundbreaking discoveries include the elucidation of conserved cell signaling and programmed cell death factors (Nobel Prize 2002), RNA interference (Nobel Prize 2006), microRNAs (Lasker Award 2008), and development of the green fluorescent protein (Nobel Prize, 2008). These discoveries and others demonstrate that this nematode can be exploited to learn about human biology in ways not possible by direct translational research. Moreover, the *Caenorhabditis* Natural Diversity Resource (CaeNDR) provides an unparalleled opportunity to investigate the genetic basis for variation in chemical exposure response and toxicity mechanisms (Crombie et al. 2023; Zdraljevic et al. 2019). Currently, CaeNDR contains 1,524 wild strains of *C. elegans* of which 1,384 have been fully sequenced, resulting in over 500 genome-wide haplotypes from distinct geographic locations (Crombie et al. 2023). These wild strains harbor genomic regions of exceptionally high genetic variation that are enriched in environmental-response genes, including xenobiotic stress-response pathways (Lee et al. 2021; Widmayer et al. 2022). Genetically diverse *C. elegans* strains exhibit variation in response to various heavy metals, pesticides, and flame retardants, demonstrating that *C. elegans* has diverse responses to a range of toxicants (Widmayer et al. 2022).

Despite its utility and tractability, only a few studies have investigated PFAS toxicity using *C. elegans*. These are limited to PFOS, PFOA, PFBA, PFBS, GenX, and a mixture of eleven PFAS chemicals (Breton et al. 2023; Feng et al. 2022; Lin et al. 2022; Sammi et al. 2019; Stylianou et al. 2019). Sammi *et al* (2019) observed that *C. elegans* exposed to levels of PFOS comparable to those detected in human blood serum induced oxidative stress and neurodegeneration, suggesting that mitochondria are a target of toxicity and pathology, as has been observed in other species (Slotkin et al. 2008; Starkov and Wallace 2002). However, specific molecular mechanisms of toxicity of all other PFAS in *C. elegans* are lacking.

The goal of this study was to characterize structure-activity relationships for PFAS chemicals and the role of natural genetic variation in response to exposures in *C. elegans*. We carefully selected thirteen PFAS chemicals based on specific structural attributes, including legacy and emerging chemicals, to test structural contributions to toxicity in an animal model (Figure 1). We measured larval growth during exposure to assess toxicity. These PFAS chemicals include PFOSA, PFBSA, PFOS, PFNA, PFOA, GenX, PFEESA, PFBS, PFPeA, PFBA, PFMOAA, 6:2 FTS and 6:2 FCA. These PFAS are among 75 that the Environmental Protection Agency prioritized for testing due to structural diversity (Patlewicz et al. 2019). Five of the PFAS in this study now have the first-ever national legally enforceable maximum contaminant levels in drinking water (PFOA, PFOS, GenX, PFNA, and mixtures of two or more of PFNA, GenX, and PFBS) (United States Environmental Protection Agengy 2024). Here, we directly compared toxicity among chemicals that vary with respect to three structural attributes, including chain length (short versus long), three different functional groups (carboxylic acids, PFCAs; sulfonic acids, PFSAs; and sulfonamides, FASAs), and chain composition (polyfluoroethers acids, PFEAs; and fluorotelomer precursors) (Figure 1). Additionally, the contribution of genetic variation to susceptibility of each PFAS was determined. Toxicity between PFAS that vary by a single structural attribute was compared among genetically diverse wild strains to identify specific structural features for which toxicity is modified by genetic variation.

**Figure 1.**
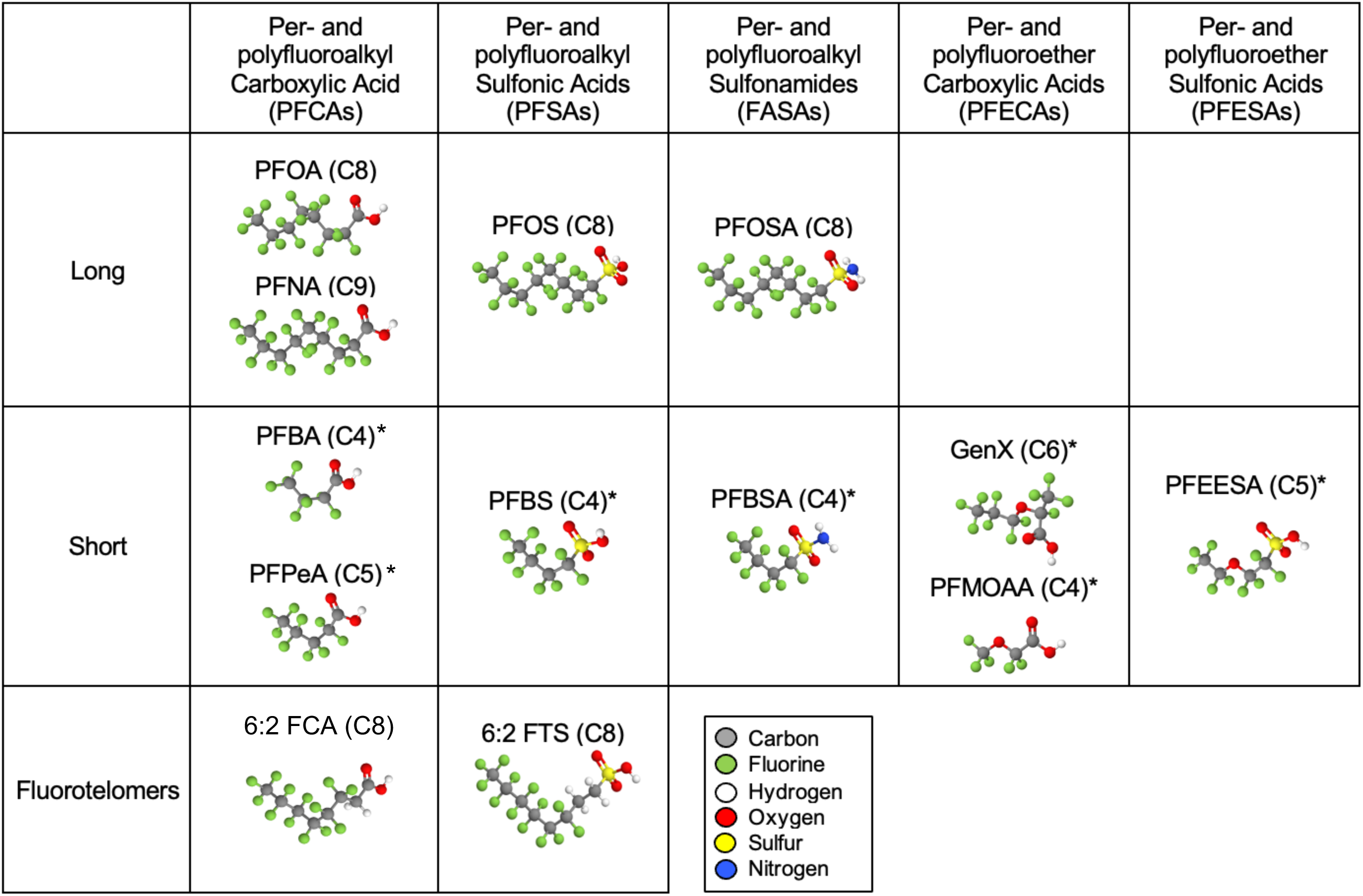
Structures of legacy and emerging per- and polyfluoroalkyl substances (PFAS) used in this study. PFAS chemicals were selected to directly compare toxicity of compounds that vary by three structural attributes: chain length (long vs short), functional group (carboxylic acid, sulfonic acid, or sulfonamides), and chain composition (polyfluoroalkyls vs polyfluoethers). Number of carbons (*eg*, 8-carbon = C8) are stated after each chemical name. Chemical structures were made using the open-source web application MolView (v2.4). *Indicates emerging PFAS chemicals.

## Methods

### C. elegans strains and maintenance

The twelve *C. elegans* isolates used in this study to measure toxicity were N2, CB4856, CX11314, DL238, ED3017, EG4725, JT11398, JU258, JU775, LKC34, MY16, and MY23. All strains were obtained from CaeNDR and comprise the “Divergent Set” (Cook et al. 2016). *C. elegans* were maintained following standard procedures at 20°C on NGM plates with *E. coli* OP50 (Brenner 1974).

### Preparation of PFAS chemicals

The following nine PFAS chemicals with respective Chemical Abstracts Service Registry Number (CASRN) were purchased from Synquest Laboratories for use in this study: perfluorooctane sulfonamide (PFOSA; 754-91-6), perfluorobutane sulfonamde (PFBSA; 30334-69-1), Ammonium perfluoro(2-methyl-3-oxahexanoate) (GenX; 62037-80-3), perfluoro(2-ethoxyethane)sulphonic acid (PFEESA; 113507-82-7), perfluorobutanesulfonic acid (PFBS; 375-73-5), perfluoropentanoic acid (PFPeA; 2706-90-3), perfluorobutanoic acid (PFBA; 375-22-4), sodium 2-perfluoromethoxyl-2,2-difluoroacetate (PFMOAA; 21837-98-9), 6:2 fluorotelomer sulfonate (6:2 FTS; 59587-39-2), and (perfluorohexyl)acetic Acid (6:2 FCA; 53826-12-3). The following three PFAS chemicals with respective CASRN were purchased from Sigma Aldrich: potassium perfluoro-1-octanesulfonate (PFOS; 2795-39-3), perfluorononanoic acid (PFNA; 375-95-1), and perfluorooctanoic acid (PFOA; 335-67-1). It is important to note that we selected the acid form of these chemicals instead of the salt form, when possible, to avoid potentially confounding results of the various salts (*e.g.* potassium, ammonium) on *C. elegans* growth. Stock solutions were prepared in methanol, except for GenX and PFMOAA, which were dissolved in sterile deionized water. Stocks were brought to neutral pH 7.0 with NaOH and stored in aliquots at −20°C. Stock solution concentrations ranged from 25 mM to 1000 mM, depending on solubility of each PFAS chemical. A fresh working solution was prepared for each experiment by thawing a single stock solution aliquot. Final dosing concentrations for each chemical used in this study are listed in Table S1.

### Chemical exposures and growth toxicity assay

All exposures were conducted in liquid cultures of fresh complete K-medium (K-plus). K-plus contains K-medium solution (32 mM KCl and 51 mM NaCl in sterile distilled water) with additions of CaCl_2_ and MgSO_4_ to a final concentration of 3 mM each, plus 5 µg/mL of cholesterol and 0.1% Ethanol (Boyd et al. 2010). To prepare each experiment, 7-10 individual L4 larvae per *C. elegans* strain were picked onto individual 10 cm NGM plates with a lawn of *E. coli* OP50. Four days later, gravid adults were hypochlorite-treated (bleached) to obtain embryos (Hibshman et al. 2021). Approximately 30 embryos from each strain were placed in eight wells of a 96-well polystyrene plate (in 50 µL of K-plus media) and placed in a 20°C shaking incubator at 180 rpm to hatch and enter L1 arrest. Additionally, ∼30 embryos of each strain were added to two wells of a separate 96-well plate to image after 24 hr to collect baseline L1 lengths (during L1 arrest) for each strain. After 24 hr, synchronized L1 larvae were dosed and fed by adding an additional 50 µL of chemical and food in K-plus. Each strain was exposed to seven doses per chemical plus appropriate vehicle control (water or methanol) (Table S1). Worms were fed a final food density of 5 mg/mL of *E. coli* HB101. This food density is sufficient to support robust growth and reproduction with this worm density, and it enables imaging the worms in liquid culture because the turbidity is relatively low. HB101 was prepared as described in Hibshman *et al*, except resuspended in K-plus media (Hibshman et al. 2021). Immediately after dosing and feeding, plates were wrapped in Parafilm to prevent evaporation and returned to the 20°C shaking incubator. 48 hr after feeding and dosing, worms were paralyzed by adding 270 µL of 50 mM sodium azide to each well and imaged after 15 minutes using a Molecular Devices ImageXpress Nano high-content imager with a 2x objective. Four independent experiments were conducted on different days with all twelve strains and all chemicals. Two preliminary experiments were conducted in just the laboratory reference strain N2 to determine the appropriate range of concentrations for each PFAS dose response curve for all twelve strains (*i.e. n* = 6 for N2).

### Image and data analysis

Images were processed on the Duke Computer Cluster via a custom CellProfiler pipeline with slight modifications (*cellprofiler-nf*; https://github.com/AndersenLab/cellprofiler-nf). Worm lengths were quantified from processed images (*easyXpress*, *R*) (Nyaanga et al. 2021; Widmayer et al. 2022). The mean length of worms per well was calculated and used for all subsequent analyses. There were no statistically significant differences between L1 lengths among strains (p = 0.28, one-way ANOVA, Figure S1), so only worm lengths greater than the minimum mean L1 starting length (206 µm) were analyzed. The *R* package ‘Analysis of Dose-Response Curves’ (*drc*) was used to estimate log_10_EC10, 20, and 50 values for each strain per treatment (Ritz et al. 2015). The ‘Lower Limit’ of the *drc* was adjusted to the minimum L1 length (206 µm). All statistical analysis and data visualization of EC50 values were conducted in RStudio. Values are conveyed as means and standard error, unless otherwise indicated. To determine variation in log_10_EC50 values, we performed ANOVA: one-way if only one factor, and two-way to determine significant PFAS-by-strain interactions, followed by Tukey’s HSD (Honestly Significant Difference) test to identify significant pairwise differences and correct for multiple comparisons (indicated in further detail in the figure legends). Broad-sense heritability values of log_10_EC50 estimates (Figure 5) and growth after exposure to mean EC50 (Figure S2) were calculated on CaeNDR (Crombie et al. 2023).

### Confirmation of Effective Concentration estimates

To validate the model estimates and approach, each strain was dosed at mean (across all strains) EC10, EC20, and EC50 values. Synchronized L1 larvae of each individual strain were exposed to the mean estimated values for each chemical and the appropriate vehicle control. Lengths were measured at 48 hr as described above, and percent growth compared to control was calculated for each strain per chemical (Figure S2). Variation in percent growth among strains was calculated for each PFAS at EC50 (Figure S3). This experiment was conducted four times. Wells were excluded from analysis if there were less than five worms per well.

### Analytical Chemistry

To determine internal body burden measurements of GenX and PFOA, synchronized N2 L4 larvae were treated with 500 *µ*M GenX, PFOA, or control conditions (0.1% MeOH) in polypropylene Erlenmeyer flasks at 5 worms/*µ*L with 25 mg/mL *E. coli* HB101. These two compounds were chosen because previous studies have compared PFOA and its replacement, GenX. After 48 h, first an aliquot of the supernatant was harvested, and then worms were washed three times after which pellets were flash frozen in liquid nitrogen and stored at −80°C. The supernatant (dosing solution) and internal body burden concentrations were determined via LC-MS/MS. The LC-MS/MS (Agilent 1260 Infinity II LC system coupled to an Agilent 6460A triple quadrupole mass spectrometer) method and sample preparation were described previously, with a few modifications for tissue extraction (Hall et al. 2022). Briefly, ∼200 mg of *C. elegans* per sample was transferred to 2 mL Eppendorf safe-lock microcentrifuge tubes. Each sample was spiked with isotopically labeled internal standards (M3GenX and M4PFOA; Wellington Laboratories; Guelph, Ontario, Canada). Next, 0.2 mL of 1 M formic acid was added, samples were vortexed for 30 seconds, then left for 5 minutes. Then, 0.8 mL of acetonitrile was added with 1 mm glass beads and homogenized in a bullet blender for 5 minutes. Homogenates were sonicated for 20 minutes without heat and centrifuged at 3,500 rpm for 5 minutes. The supernatant was transferred to a 15 mL polypropylene centrifuge tube. The pellet was resuspended in 1.0 mL of acetonitrile to repeat the extraction process, and supernatants were combined. The supernatant was centrifuged for 10 minutes at 3,500 rpm and transferred to a new 15 mL tube. Extracts were concentrated to near dryness under nitrogen at 35°C, then resuspended in 1000 µL of 1:1 methanol:water, filtered through a 0.2 µm nylon syringe filter into a LC vial. Prior to LC-MS/MS analysis, an isotopically labeled standard (M2PFOA) was spiked into samples to assess recovery of internal standards. Recovery averaged 150% (71 to 204%) for GenX and 61% (56 to 65%) for PFOA. Bioconcentration factor (BCF) results were calculated as [worm]/[supernatant] for GenX or PFOA (Table S4).

### Data availability

Raw data and code used for data analyses and visualization are available at https://github.com/tessleuthner/2024-PFAS-DS.

## Results

### Effects of PFAS chemicals on N2 growth

To determine the effects of PFAS exposure on *C. elegans* growth, N2 larvae synchronized by L1 arrest were exposed to a range of concentrations of PFOSA, PFBSA, PFOS, PFNA, PFOA, GenX, PFEESA, PFBS, PFPeA, PFBA, 6:2 FTS, 6:2 FCA, and PFMOAA (Figure 2 and Table S1). After 48 hr of exposure during larval development, images were acquired and analyzed to extract worm length (Figure 2C-E). Lengths at each dose were used to calculate the concentration that caused 50% growth inhibition (EC50) for each chemical (Figure 2F, Table 1). We were unable to estimate EC50 values for three of the chemicals (PFMOAA, 6:2 FTS, and 6:2 FCA), as these PFAS were insoluble at concentrations that sufficiently inhibited growth (Figure 2E). Therefore, these three chemicals were omitted from further experiments throughout this study.

**Figure 2.**
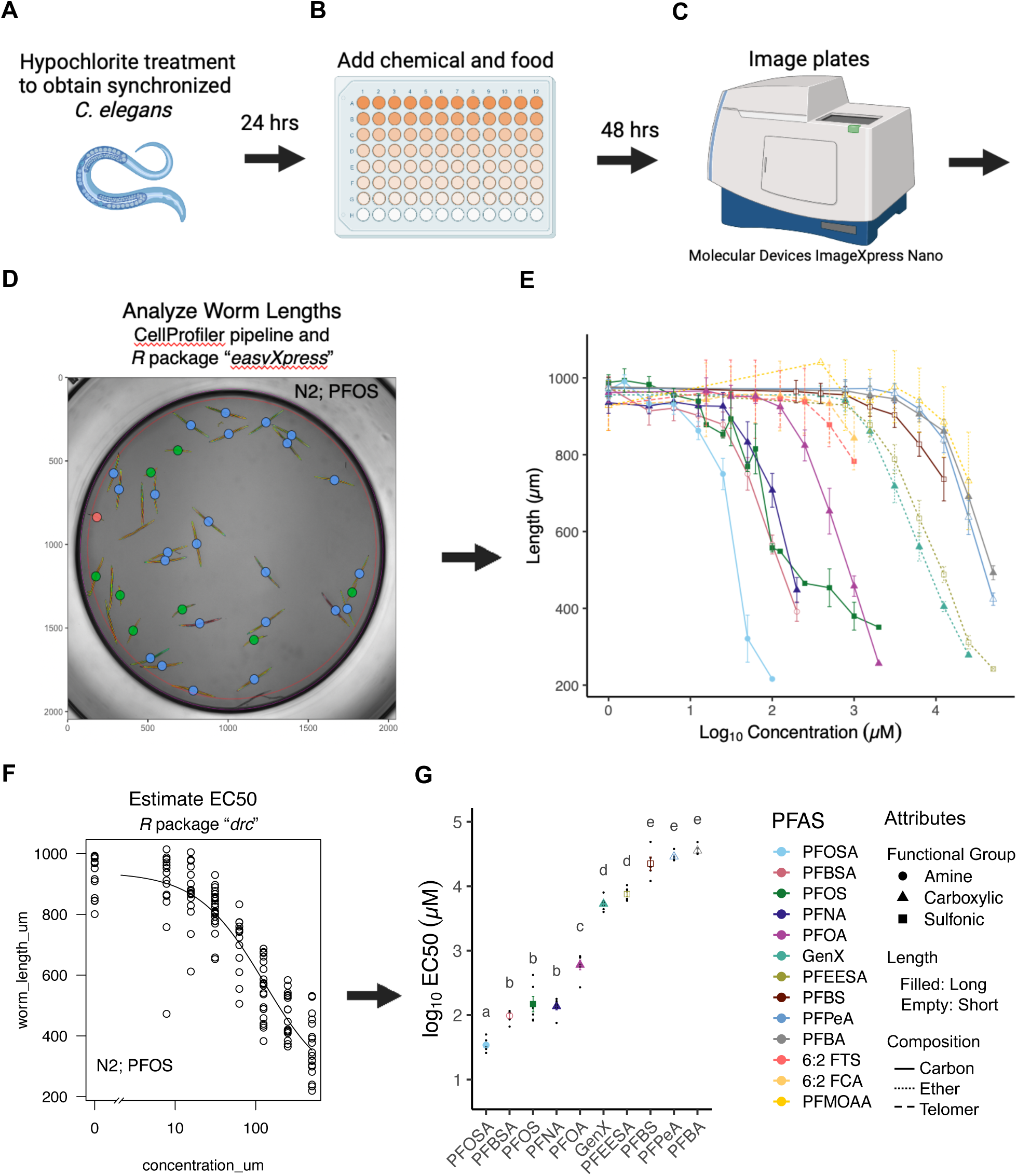
Experimental design and the effects of thirteen PFAS chemicals on growth in the laboratory-adapted, wild-type reference strain, N2. (A) Gravid adults were hypochlorite-treated to obtain embryos, and a target of 30 embryos were placed in each well of a 96-well plate. (B) 24 hr after bleach, once the embryos had hatched and entered L1 arrest, food and chemical were added to synchronized L1s to initiate larval development and PFAS exposure. (C) Plates green; and L1, red). Worm lengths were obtained using the *R* package *easyXpress*. (E) Mean worm length per well per dose was calculated. The legend is below in panel G. (F) To estimate concentrations in which there was 50% growth inhibition compared to control (EC50), the *R* function *drc* was used to estimate an EC50 value for each experimental replicate. The plot shown is from one experimental replicate in which N2 larvae were exposed to seven concentrations of PFOS for 48 hr, plus control. Each empty dot is an individual worm length. (G) EC50 values were log-transformed to determine variation in log_10_EC50 among ten PFAS chemicals in N2 (one-way ANOVA, *p* < 1e^−40^; letters indicate groups of chemicals that are not significantly different from each other based on Tukey HSD post-hoc analysis, including correction for multiple comparisons). Solubility limited the maximum exposure concentration of the chemicals 6:2 FTS, 6:2 FCA, and PFMOAA; therefore, EC50 values could not be calculated. Dots represent the average log_10_EC50 of six experimental replicates. Error bars are standard error of the mean. Color, shape, and fill correspond to chemical, functional group, and the chain length respectively. Lines indicate the chain composition. (A-C) Created with BioRender.com.

**Table 1.** EC50 (µM) of 10 PFAS chemicals in wild-type and among twelve *C. elegans* strains. The mean, minimum, maximum, and range of EC50 between the twelve strains was calculated for each PFAS. The variation in toxicity between strains was determined between the most sensitive and the most resistant strain within each PFAS.

| Chemical | N2 EC50 ($\mu\text{M}$)<br>(mean ± SD) | Divergent Set EC50<br>($\mu\text{M}$) (mean ± SD) | Minimum<br>EC50 ($\mu\text{M}$) | Maximum<br>EC50 ($\mu\text{M}$) | Range of<br>EC50 ($\mu\text{M}$) | Fold-change<br>toxicity |
| --- | --- | --- | --- | --- | --- | --- |
| PFOSA | 35.0 (± 8.79) | 35.9 (± 8.38) | 27.9 | 41.4 | 13.6 | 1.5 |
| PFBSA | 99.9 (± 19.8) | 81.1 (± 4.09) | 65.2 | 104.0 | 39.1 | 1.6 |
| PFNA | 142.0 (± 36.2) | 138.0 (± 8.68) | 80.0 | 171.0 | 91.3 | 2.1 |
| PFOS | 182.0 (± 137) | 115.0 (± 20.6) | 95.9 | 155.0 | 58.6 | 1.6 |
| PFOA | 640.0 (± 210) | 746.0 (± 103) | 482.0 | 845.0 | 363.0 | 1.8 |
| GenX | 5,480.0 (± 1,420) | 5,790.0 (± 395) | 3,950. | 8,860.0 | 4,920.0 | 2.2 |
| PFEESA | 7,710.0 (± 1,790) | 9,920.0 (± 395) | 7,360.0 | 16,600.0 | 9,270.0 | 2.3 |
| PFBS | 25,200.0 (± 14,400) | 26,400.0 (± 7,330) | 17,800.0 | 35,300.0 | 17,500.0 | 2.0 |
| PFPeA | 29,100.0 (± 4,490) | 31,500.0 (± 627) | 27,300.0 | 40,100.0 | 12,800.0 | 1.5 |
| PFBA | 36,100.0 (± 6,530) | 36,900.0 (± 1,590) | 31,900.0 | 49,900.0 | 18,000.0 | 1.6 |

The log_10_EC50 values for the ten PFAS chemicals varied substantially (one-way ANOVA, *p* < 1e^−40^, Figure 2G). The least toxic chemical, PFBA, was more than 1000-fold less toxic than the most toxic chemical, PFOSA (*p* < 1e^−13^). The long-chain sulfonamide, PFOSA (C8), had the greatest toxicity with a mean EC50 of 35 µM (± 8.79). The short-chain sulfonamide, PFBSA (C4) was less toxic than its long-chain complement, with a mean EC50 of 99.9 µM (± 19.8) (*p* < 1e^−3^). PFBSA had similar toxicity to the long-chain carboxylic acid PFNA (C9), which had a mean EC50 of 142 µM (± 36.2) (*p* = 0.82), and the long-chain sulfonic acid PFOS (C8), which had a mean EC50 of 182 µM (± 137) (*p* = 0.58). There was no difference in toxicity between PFNA and PFOS (*p* = 1). The next most toxic PFAS was PFOA (C8), which had a mean EC50 value of 640 µM (± 210) and was 3.5-fold less toxic than PFOS (*p* < 1e^−6^). PFOA has one fewer carbon than PFNA and was more than 4-fold less toxic (*p* < 1e^−6^). The two perfluorether acids, GenX (carboxylic acid) and PFEESA (sulfonic acid), had similar toxicity with EC50 values of 5,480 µM (± 1,416) and 7,710 µM (± 1,790), respectively (*p* = 0.8). GenX was more than eight times less toxic than the legacy chemical that it replaced, PFOA (*p* < 1e^−13^), despite a similar bioconcentration factor (Figure S4). PFEESA had similar toxicity to GenX. PFEESA was about 3-fold more toxic than the next most toxic chemical, a short-chain sulfonic acid, PFBS (C4), which had an EC50 value of 25,200 µM (± 14,400) (*p* < 1e^−3^). PFBS was similar in toxicity to both short-chain carboxylic acids PFPeA (C5) and PFBA (C4), which had EC50 values of 29,100 µM (± 4,490) and 36,100 (± 6,530), respectively (*p* = 0.97, 0.48). The two short-chain carboxylic acids, PFPeA and PFBA, had similar toxicity (*p* = 1).

### Variation in EC50 among twelve genetically diverse C. elegans strains

To determine effects of PFAS on growth in genetically diverse strains, synchronized L1 larvae from twelve genetically diverse strains were exposed for 48 hr to the same range of concentrations of PFOSA, PFBSA, PFOS, PFNA, PFOA, GenX, PFEESA, PFBS, PFPeA, and PFBA (Figure 3 and Table S1). The EC50 value was estimated for each chemical and each strain, and the average value across all strains was calculated for each chemical (Figure 4A, Table 1). The EC50 of the most sensitive and resistant strains within each chemical were also calculated (Table 1). Similar to N2 results, the mean EC50 values for the 10 PFAS chemicals varied by three orders of magnitude (one-way ANOVA, *p* < 1e^−40^) and share toxicity rankings. The long-chain sulfonamide, PFOSA (C8), had the greatest toxicity, with a mean EC50 of 35.9 µM (± 8.38). The short-chain sulfonamide, PFBSA (C4) was less toxic than its long-chain complement, with a mean EC50 of 81.1 µM (± 4.09) (*p* < 1e^−7^). PFBSA was no more toxic than PFOS (C8), which had a mean EC50 of 115.0 µM (± 20.6) (*p* = 0.066). The next most toxic PFAS was PFNA (C9) which had a mean EC50 of 138.0 µM (± 8.68) and was similar to PFOS (*p* = 0.5). There was more than a 5-fold increase in EC50 value in the next most toxic PFAS, PFOA (C8), which had a mean EC50 value of 746.0 µM (± 103). The two perfluoroether acids GenX and PFEESA, which had EC50 values of 5,790 µM (± 395) and 9,920 (± 395) µM, respectively, were significantly less toxic than PFOA. Consistent with N2 results, GenX was roughly eight times less toxic PFOA (*p* < 1e^−13^). Among the twelve strains, PFEESA was about half as toxic as GenX (*p* < 0.001). The next most toxic PFAS was the short-chain sulfonic acid, PFBS (C4), with an EC50 value of 26,400 µM (± 7,330). PFBS was similar in toxicity to the short-chain carboxylic acid of similar length, PFPeA (C5), which had an EC50 of 31,530 µM (± 627) (*p* = 0.31), though PFBS was more toxic than the other short-chain carboxylic acid, PFBA (C4), which had an EC50 of 36,900 (± 1,600) (*p* < 0.05). Both short-chain carboxylic acids, PFPeA and PFBA, had similar toxicity (*p* = 0.85). Each EC50 estimate per PFAS chemical was confirmed by exposing each individual strain to the mean estimated EC50 value (across all strains) for 48 hr and quantifying percent growth (Figure S2). Estimated EC10 and EC20 values were also tested for each chemical.

**Figure 3.**
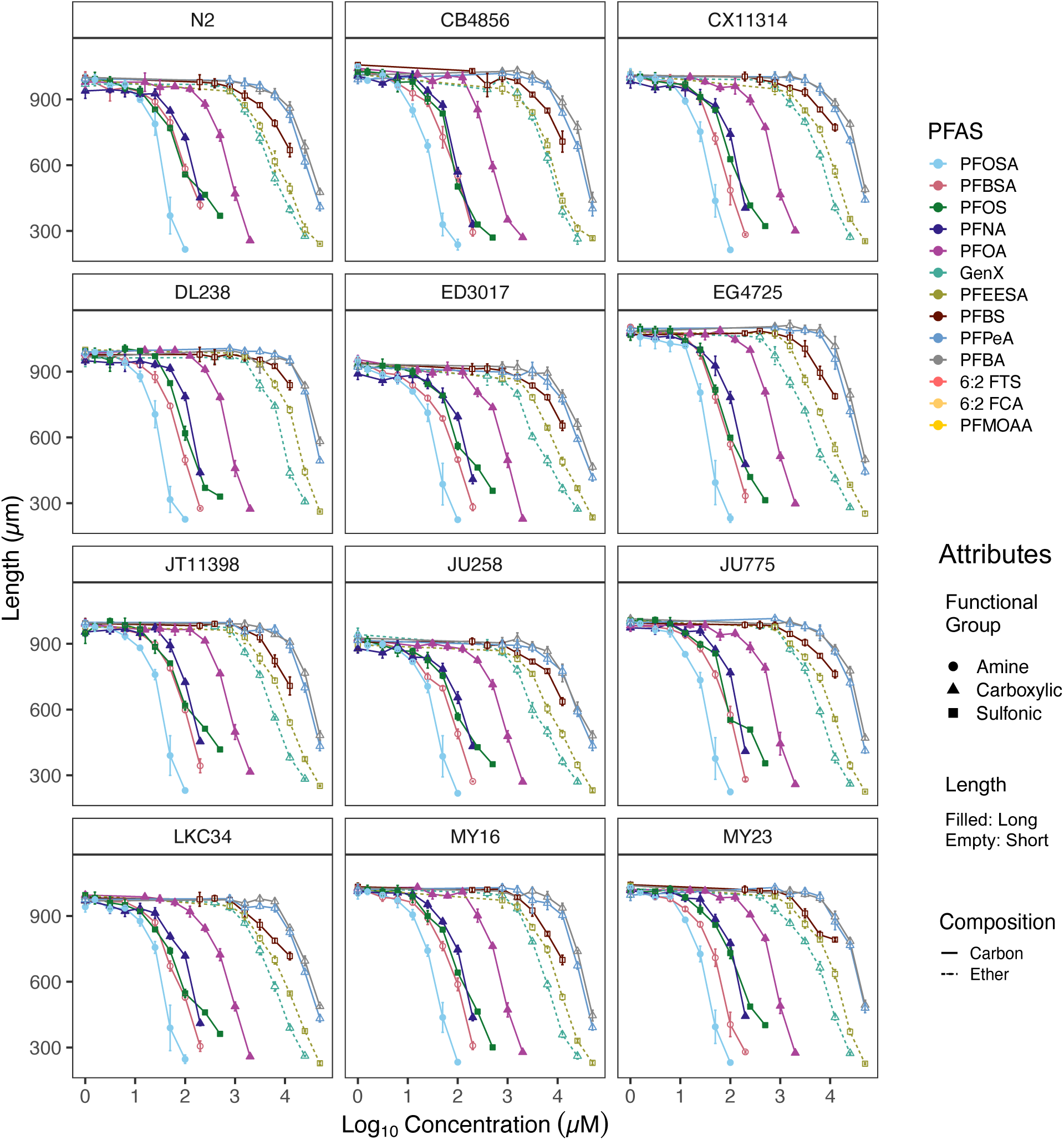
Effects of 48 hr larval PFAS exposure on size of twelve *C. elegans* strains. Worm length was measured in the wild-type, laboratory-adapted reference strain, N2, and eleven wild *C. elegans* strains after 48 hr of post-embryonic exposure to various concentrations of ten PFAS chemicals. Mean worm length was calculated for each concentration of each chemical per replicate (an average of 18 ± 6 (standard deviation) individuals/well). Dots represents the average length of four experimental replicates. Error bars are standard error of the mean. Color, shape, and fill correspond to chemical, functional group, and chain length, respectively. Lines indicate chain composition. Dose-response curves could not be fit to three PFAS (6:2 FTS, 6:2

**Figure 4.**
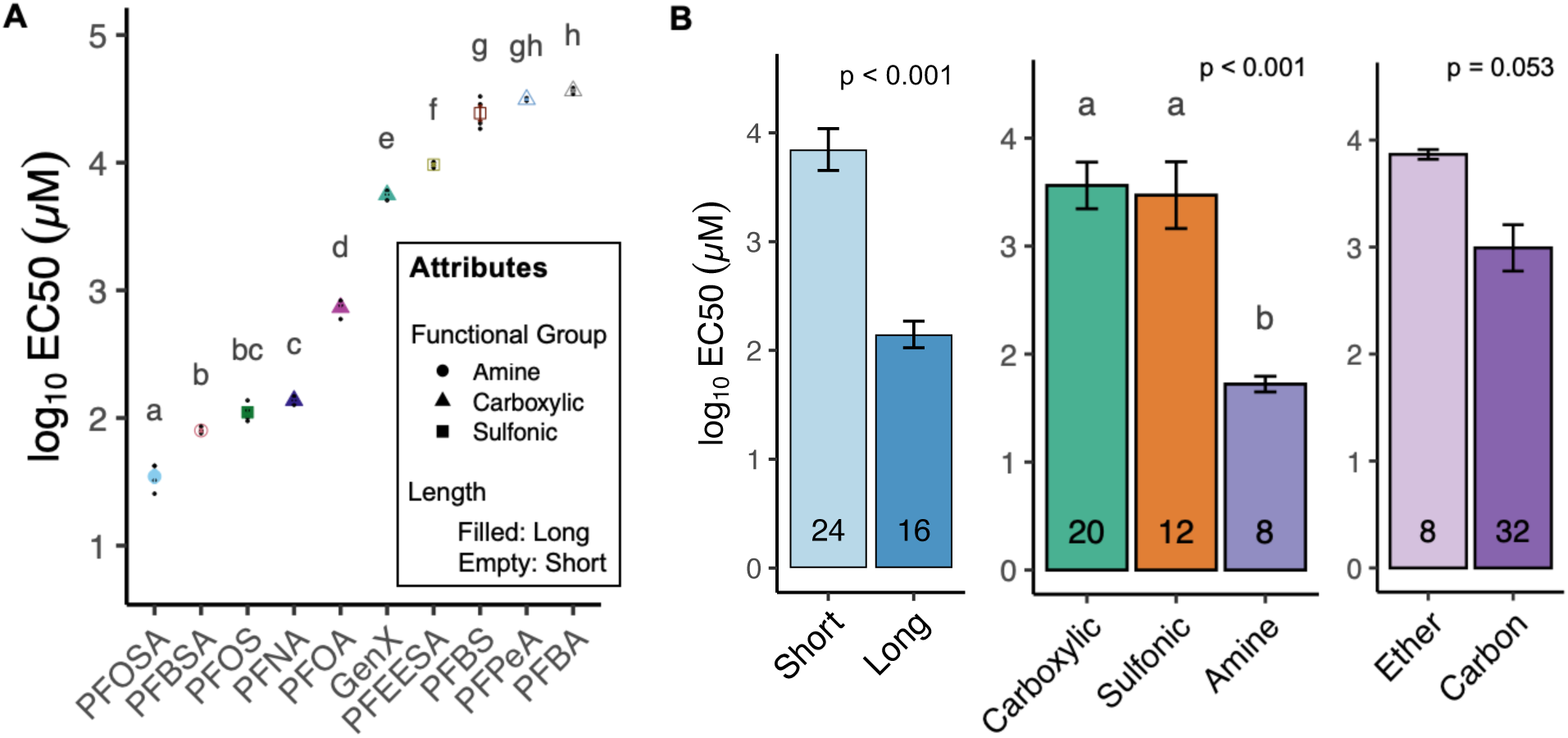
Variation in EC50 among PFAS chemicals and contribution of structural attributes to toxicity. (A) Dose-response curves were fit for each individual strain. Effective concentrations which inhibited growth by 50% (EC50) were calculated and log_10_ transformed. The mean EC50 for each PFAS chemical across all twelve strains was calculated. Each colored point represents the mean EC50 for four experimental replicates. Small black dots are mean EC50 values for each experimental replicate. Chemicals are in rank order of greatest to lowest toxicity (lowest to highest EC50 values). EC50 values are statistically different among chemicals (one-way ANOVA, *n* = 4, *p* < 1e^−40^). Different letters indicate significantly different EC values after correction for multiple comparisons (Tukey HSD). Error bars indicate standard error of the mean. Color, shape, and fill correspond to chemical, functional group, and the chain length respectively. (B) The mean log_10_EC50 value across the twelve strains was determined for each PFAS that have each indicated structural attribute. The number on each bar indicates N, or the total number of PFAS chemicals analyzed multiplied by four experimental replicates (*p*-value is from one-way ANOVA; different letters indicate significant differences after correction for multiple comparisons (Tukey HSD)).

### Toxicity of PFAS that vary by structural attributes

To determine effects of chain length, functional group, and presence of an ether (chain composition) on toxicity, we compared log_10_EC50 values between chemicals containing each molecular attribute (Figure 4B). Long-chain PFAS (PFOA, PFNA, PFOS, and PFOSA) had a mean log_10_EC50 of 2.15 and were on average 50 times more toxic than the short-chain PFAS (PFBA, PFPeA, PFBS, PFBSA, GenX, and PFEESA) which had a mean log_10_EC50 of 3.85 (one-way ANOVA, *p* < 0.001). There was significant variation in toxicity due to functional group independent of chain length or composition (*p* < 0.001). The perfluoroalkyl sulfonamides (FASAs) had a mean log_10_EC50 of 1.72 and were significantly more toxic than the carboxylic acids (PFCAs), and the perfluoroether carboxylic acid GenX, which had a mean log_10_EC50 of 3.56 (*p* < 0.0001). The FASAs were also significantly more toxic than the perfluoroalkyl sulfonic acids (PFSAs), and the perfluoroether sulfonic acid PFEESA, which had a mean log_10_EC50 of 3.47 (*p* < 0.001). There was no difference in toxicity between PFAS with a carboxylic acid and sulfonic acid functional group (*p* = 0.96). To determine the contribution of chain composition to toxicity, we compared log_10_EC50 values of the two perfluorethers GenX and PFEESA to those without an ether, independent of chain length and functional group. PFAS with ethers had a mean log_10_EC50 of 3.87, while those without had a lower mean log_10_EC50 of 2.99, though not statistically more toxic (*p* = 0.053).

### Natural variation in PFAS toxicity among diverse C. elegans strains

We determined variation in toxicity by comparing log_10_EC50 values among the twelve genetically diverse *C. elegans* strains within each chemical (Figure 5). If variation among strains for a given chemical was statistically significant, strain-specific interactions were determined to investigate variation in susceptibility between individual strains (Table S2). Broad-sense heritability estimates (*H^2^*) were calculated to estimate the contribution of genetic variation to variation in response to exposure to each PFAS (Figure 5; Cook et al. 2016). The *H^2^* proportion ranges from 0 to 1, where higher estimates indicate a greater contribution of genetics to phenotypic variation.

**Figure 5.**
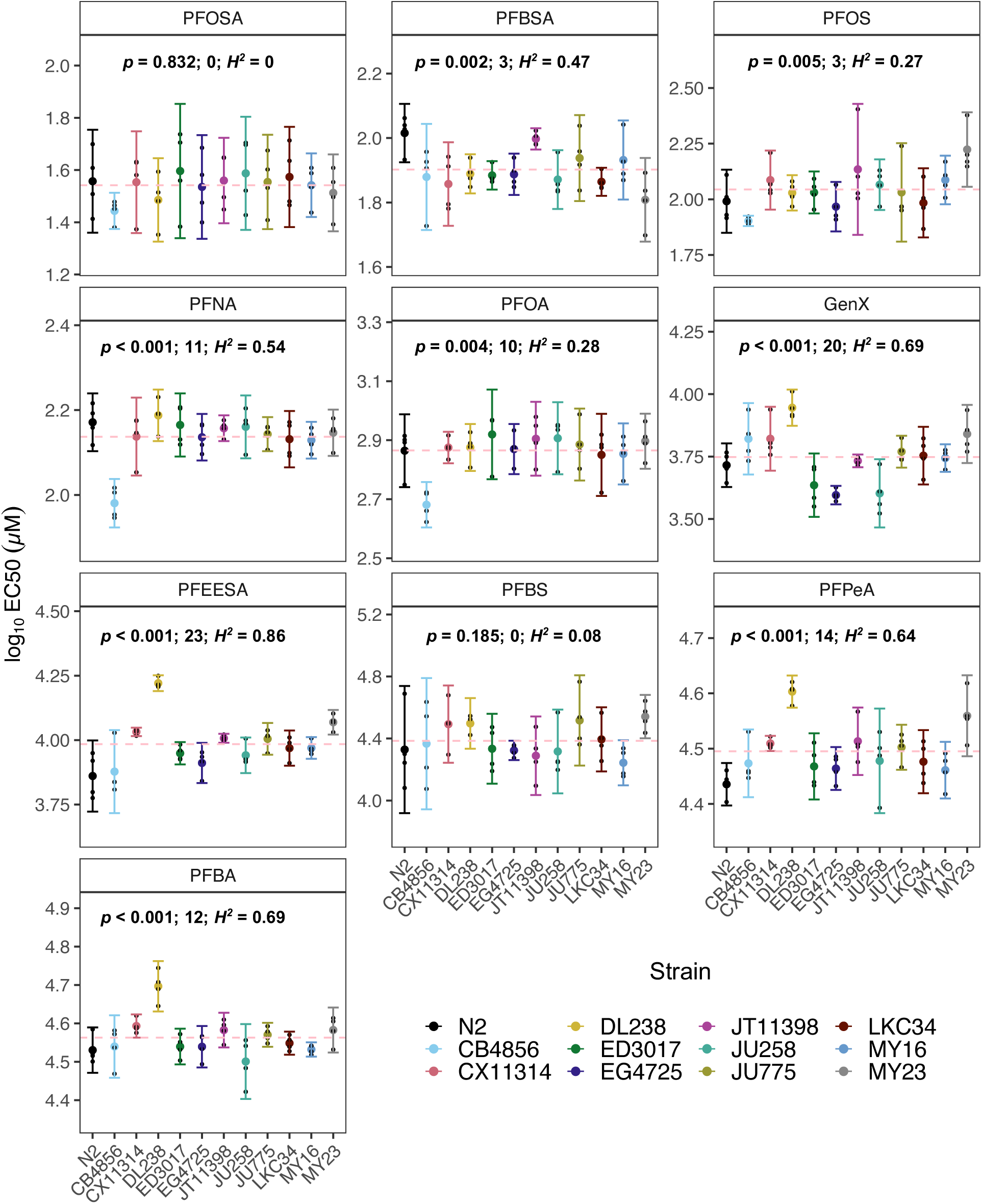
Variation in PFAS toxicity across twelve wild *C. elegans* strains. Dose-response curve models were fit for each strain to estimate strain-specific EC50 values (log_10_-transformed) for each experimental replicate (*n* = 4). Each colored dot represents the mean EC50 and small black dots indicate log_10_EC50 for each experimental replicate. The red dashed line is the mean overall significant strain*treatment interaction (*p* < 1.41e^−9^, two-way ANOVA) among all strains and PFAS chemicals. Differences in log_10_EC values among strains were then determined within each chemical (*p*-value, one-way ANOVA). If there was significant variation detected, within-treatment differences in log_10_EC values between each pair of strains were evaluated for significance using Tukey’s HSD to identify significant pairwise differences and correct for multiple comparisons. The one-way ANOVA *p-*value and number of significant pairwise differences are displayed. The specific pairwise differences that are significant are listed in Table S2. Broad-sense heritability was estimated for each log_10_EC value as the trait (*H^2^*) and is also displayed.

PFBSA, PFOS, PFNA, PFOA, GenX, PFEESA, PFPeA, and PFBA all had significantly different log_10_EC50 values across the twelve strains (Figure 5). When exposed at the mean EC50 values, there was also significant variation in growth among strains after exposure to PFNA, GenX, PFBS, PFPeA, and PFBA (Figure S2, Table S3). Among these PFAS chemicals, there was a range in the number of significant differences between strains and the broad-sense heritability values. Response to exposure to PFEESA resulted in 23 significant pairwise strain interactions and the highest broad-sense heritability value (*p* < 0.001, *H^2^ =* 0.86). The laboratory strain, N2, was significantly different from five other strains, while the most resistant strain, DL238, had a significantly higher log_10_EC50 compared to all 11 other strains exposed to PFEESA. In fact, among three other PFAS chemicals (GenX, PFBA, and PFPeA), DL238 also had the highest log_10_EC50 values. Response to GenX was also highly variable among strains as there were 20 significant differences among strains in response to GenX exposure and *H^2^ =* 0.69 (*p* < 0.001). PFBA also had a high heritability value of 0.69, with twelve significant differences among strains (*p* < 0.001). This variation is driven by the high log_10_EC50 of DL238, which was different from all eleven other strains. The other short-chain PFCA used in this study, PFPeA, had the next highest heritability value (*H^2^ =* 0.64) with fourteen significant differences among strains (*p* < 0.001). DL238 had a significantly higher log_10_EC50 value than ten other strains (all except MY23). Unlike the short-chain carboxylic acids, the short-chain sulfonic acid, PFBS, did not have significant variation in log_10_EC50 among strains, and it had a very low heritability value of *H^2^ =* 0.08 (*p* = 0.12). However, the long-chain sulfonic acid, PFOS, did have significant variation among strains (*p* < 0.001). The only differences in response to PFOS exposure were between MY23 (which had the highest log_10_EC50) compared to CB4856, EG4725, and LKC34. PFOS also had lower heritability (*H^2^ =* 0.27). The long-chain carboxylic acids PFNA and PFOA had significant variation in strain response (*p* < 0.001 and *p* < 0.01, respectively). In both cases, variation was driven by the strain CB4856. CB4856 had the lowest log_10_EC50 compared to all eleven other strains after exposure to PFNA, and the lowest compared to all strains except LKC34 after exposure to PFOA. There was no significant variation among strains that were exposed to the most toxic PFAS in this study, PFOSA (*p* = 0.83). However, exposure to the short-chain sulfonamide, PFBSA, did have significant differences in log_10_EC50 among strains, but only between the strain with the highest log_10_EC50 value, N2, and the two lowest log_10_EC50 values, CX11314 and MY23 (*p* < 0.01). PFBSA had a moderate heritability value of *H^2^ =* 0.47.

### PFAS structure-specific variation in strain susceptibility

Analysis of effects on growth within each individual PFAS chemical revealed that there was variation among strains in response to exposures to eight of the ten PFAS studied (Figure 5). However, we were interested in comparing the variation in toxicity among strains between chemicals with differences in specific structural attributes. To determine the contribution of genetic variation to structure-specific responses, we compared the log_10_EC50 values between chemicals that differ by one specific structural attribute (*n* = 4 replicates per strain per treatment, two-way ANOVA). Statistical comparisons reported were conducted on log_10_EC50, but for better data visualization we plotted the average mean-normalized values per strain (Figure 6). Comparisons between chemicals that did not exhibit significant variation among strains are displayed in Figure S5. Overall, when we examined pairwise comparisons between PFAS that vary by one structural attribute, we observed variation among the strain response in eight of the nineteen possible comparisons.

**Figure 6.**
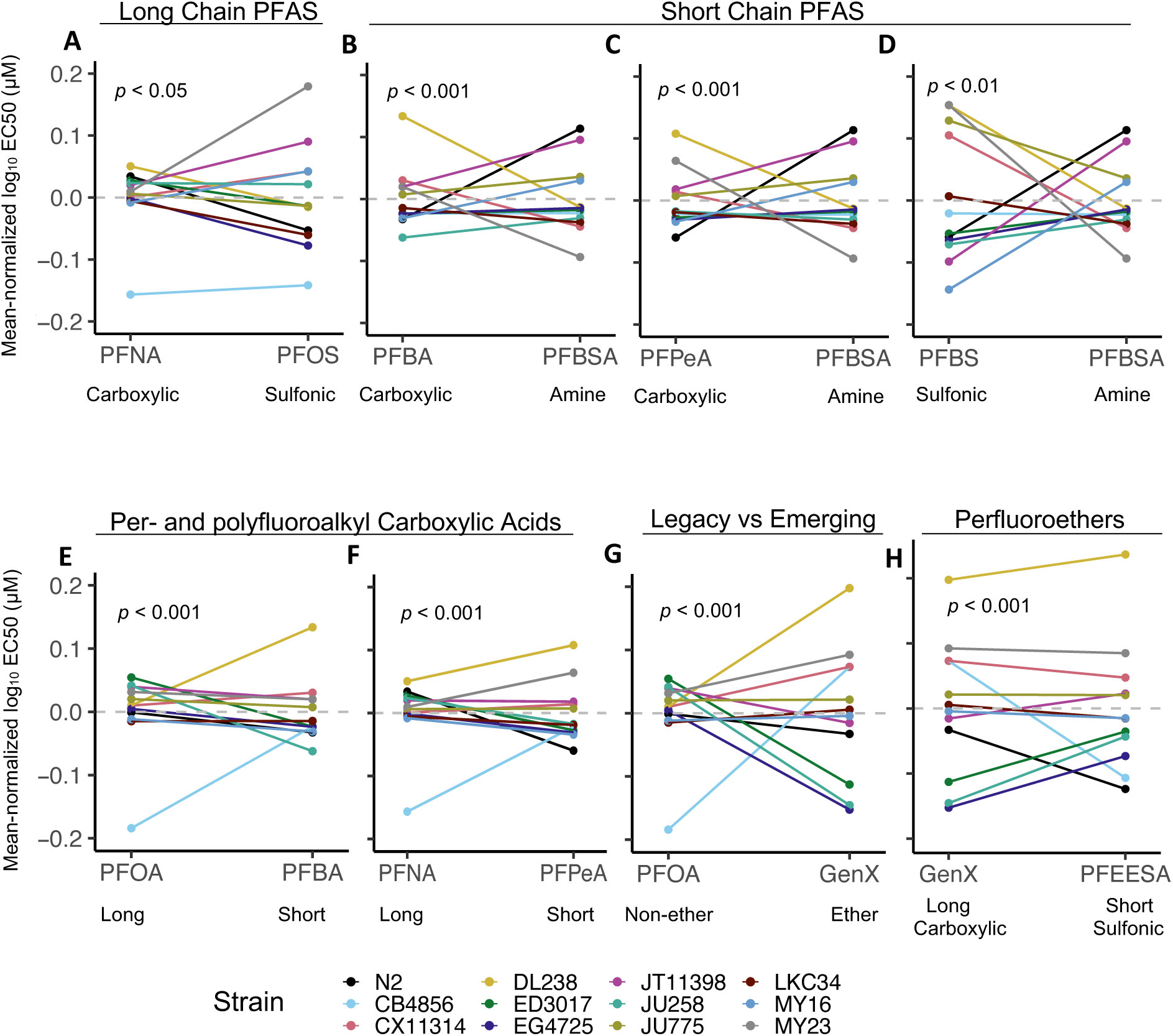
Strain and structure-specific variation in susceptibility to PFAS chemicals. Variation in toxicity (log_10_EC50 per chemical) among strains between specfic PFAS comparisons was determined (*p*-value is from two-way ANOVA, PFAS*strain interaction). EC50 values were mean-normalized and plotted to better visualize variation in strain response between each PFAS pair. Variation among strains in response to functional group within (A) long-chain PFAS (carboxylic acid vs sulfonic acid) and (B-D) short chain PFAS (carboxylic acid and sulfonic acids compared to sulfonamide). (E, F) Variation in strain response between long- and short-chain PFCAs. (G) Variation in strain response between PFOA (legacy, non-ether) and GenX (emerging, ether). (H) Variation in strain response between two ethers (GenX and PFEESA) that vary by both length and functional group. Each strain is represented by the same color in each panel. Each dot represents the mean of each strain mean-normalized log_10_EC50 (µM) (*n* = 4 experimental replicates per strain per treatment). All raw log_10_EC50 comparisons for PFAS that vary in a single structural attribute (including insignificant comparisions) are shown in Figure S5.

We investigated natural variation in susceptibility between all pairs of long-chain PFAS chemicals that vary by functional group. There was significant variation among strains when we compared PFNA to PFOS (Figure 6A, *p* < 0.05). However, there was no significant variation in strain response when we compared PFOA to PFOS or PFNA (Figure S5A, B, *p* = 0.26 and *p* = 0.97, respectively). We also observed no variation among strain response when we compared PFOA and PFOSA (Figure S5C, *p* = 0.96), PFOS and PFOSA (Figure S5D, *p* = 0.24), and PFNA and PFOSA (Figure S5F, *p* = 0.90).

Next, we investigated natural variation in susceptibility between functional groups within the short-chain PFAS. There was significant variation among strains when we compared PFBA, PFPeA, and PFBS to PFBSA (Figure 6B, C, and D; *p* < 0.001, *p* < 0.001, and *p* < 0.01, respectively). There was no significant variation among strains between the two PFCAs, PFBA and PFPeA (Figure S5G, *p* = 0.59). There was also no significant variation among strains when we compared PFBA to PFBS, nor PFPeA to PFBS (Figure S5H and K, *p* = 0.52 and 0.61, respectively).

We next determined the effect of chain length on natural variation in strain response. There was significant variation in strain response between long versus short carboxylic acids when we compared PFNA (C9) to PFPeA (C5), and PFOA (C8) to PFBA (C4) (Figure 6E, F; *p* < 0.001 and *p* < 0.001, respectively). However, there was no significant variation in response among strains between long and short-chain sulfonic acids or sulfonamides (Figure S5O and P, *p* = 0.18 and *p* = 0.43, respectively).

To investigate variation in response among strains to PFAS that varied by chain composition, we compared two long-chain PFAS chemicals: the legacy chemical, PFOA (no ether) with its replacement, GenX (ether). There was significant variation among strains in response to these two chemicals (Figure 6G, *p* < 0.001). There was no significant variation among strain response when we compared the two short-chain PFAS chemicals PFBS (no ether) and PFEESA (ether) (Figure S5Q, *p* = 0.45). We did observe variation in strain response to the two ethers, GenX and PFEESA, though it is important to note that these chemicals differ by both chain length and functional group (Figure 6H, *p* < 0.001).

## Discussion

The role of genetic variation in the response to exposure to environmental pollutants in diverse populations is largely unknown. However, this is critical for assessing risk, as environmental pollutants are the leading cause of premature death globally (Landrigan et al. 2018), and humans are genetically diverse. To our knowledge, we provide the first evidence that genetic variation contributes to susceptibility to PFAS toxicity. PFAS pollution is ubiquitous, and exposure is associated with many diseases such as various cancers, neurodegenerative disorders, and harmful effects on the reproductive, immune, and endocrine systems. Unfortunately, there are currently more than 14,500 unique PFAS chemicals identified of undetermined toxicity (EPA CompTox), and interindividual variation in susceptibility and health consequences are unknown. Our analysis suggests that genetic variation not only modifies susceptibility to PFAS exposure but may affect structure-specific toxicity. This variation in toxicity will be a useful tool to investigate molecular mechanisms of structure-specific toxicity.

It is impractical to individually test each PFAS chemical, so we carefully selected PFAS that vary in chain length, functional group, and chain type (ether vs non-ether), including legacy and emerging PFAS. We conducted experiments to investigate the effects of these specific structural attributes on growth in the nematode *C. elegans* to elucidate structure-activity relationships (Figure 1, 2). We harnessed the natural genetic diversity among the reference strain, N2, and eleven genetically diverse, wild strains of *C. elegans* to determine the contribution of genetic variation to toxicity across PFAS and with respect to specific PFAS structural features (Figure 3 and 4). Our results demonstrate that the variation in PFAS toxicity across ten diverse chemicals is largely driven by chain length and functional group. This is consistent with previous studies that demonstrate long-chain PFAS are more potent than short-chain PFAS, and the perfluorosulfonamides (FASAs) are more toxic compared to the perfluoroalkyl carboxylic acids (PFCAs) or perfluoroalkyl sulfonic acids (PFSAs) (Figure 4). Across the PFAS investigated in this study, we found that an astounding 8-86% of phenotypic variation among strains was due to genetic variation (broad-sense heritability) among nine of the ten PFAS, and that toxicity varied by 1.5 to 2.3-fold between the most sensitive and resistant strains (Figure 5, Table 1). Furthermore, we observed variation among *C. elegans* strains in their susceptibility to some specific PFAS structures (Figure 6). Our results suggest that there are structure-specific molecular mechanisms of PFAS toxicity to investigate further, which could help to better triage PFAS for toxicity testing and risk assessment. This study reveals the importance of including genetically diverse organisms in toxicological studies to understand variation in susceptibility to PFAS exposures, and we demonstrate that *C. elegans* provides a powerful study system for doing so.

The order of PFAS toxicity across the ten chemicals used in this study in the reference *C. elegans* strain, N2, was PFOSA > PFBSA ≈ PFNA ≈ PFOS > PFOA > GenX ≈ PFEESA > PFBS ≈ PFPeA ≈ PFBA (Figure 2G). This order of potency was similar to the average EC50 values of 12 genetically diverse strains used in this study. Including diverse strains did change the rank order of PFOS and PFNA, which is notable, but not statistically significant (Figure 4A). Our identification of the sulfonamide PFOSA (C8) as the most toxic PFAS agrees with previous studies that also screened multiple PFAS chemicals in larval zebrafish and *in vitro* (Dasgupta et al. 2020; Slotkin et al. 2008; Truong et al. 2022). The high potency of PFOSA may be partially attributed to the highly reactive functional head group, and because PFOSA induces oxidative stress, is a mitochondrial uncoupler, and even inhibits DNA replication *in vitro* (Slotkin et al. 2008; Starkov and Wallace 2002). This likely contributes to perturbation in lipid metabolism and developmental neurotoxicity observed after exposure to PFOSA in these *in vitro* and *in vivo* studies. PFOSA is a precursor to PFOS, and biotransformation may occur via glucuronidation mechanisms (Xie et al. 2009; Ross et al. 2012). To our knowledge, there is no consensus on which metabolic derivative contributes to specific mechanisms of action of PFOSA. This suggests that variation in UDP glucuronyltrasnferase (UDPGT) activity between species, strains, or even individuals within a population could result in variation in PFOSA metabolites and thus variation in toxicity (Martin et al. 2010). Though we observed no variation among the twelve genetically diverse strains of *C. elegans* in response to PFOSA, we did observe variation in sensitivity to PFOS among three strains (Figure 5). The transformation of PFOSA or any precursor in *C. elegans* is yet to be investigated and is an exciting potential future direction. PFOS has been phased out of production, but the long half-life of PFOSA in the environment could contribute to high blood serum levels of PFOS in humans and in tissues sampled from wildlife. Furthermore, the precursor to PFOSA (EtPFOSA) is the active ingredient in Sulfluramid®, one of the most commonly used insecticides in Brazil and parts of Latin America, which renders this a major health concern (Nascimento et al. 2018; Zhang et al. 2021).

We found that the sulfonamide functional head group drives toxicity, as the next most toxic PFAS is the short-chain counterpart to PFOSA, PFBSA (Figure 4B). PFBSA is likely an emerging PFAS as it was found to be the major metabolite of post-2002 Scotchgard fabric protector (3M Company) in rat liver microsomes (Chu and Letcher 2014). This emerging FASA has been detected in tissues of multiple fish species, in addition to the blood and serum of cattle that were exposed to AFFF-contaminated groundwater (Chu et al. 2016; Dewapriya et al. 2023). However, very few studies have investigated the potential toxicity and adverse health effects of PFBSA. Similar to this study, Rericha *et al*. discovered that PFBSA was significantly more toxic than three other PFAS of the same length that varied only by functional head group: PFBS, (sulfonate), 4:2 FTS (sulfonate), and PFPeA (carboxylate) (Rericha et al. 2022). Unlike PFOSA, we do observe variation among strains in response to PFBSA (Figure 5).

The next most potent chemicals in this study are the three other long-chain PFAS: PFOS, PFNA, and PFOA (Figure 2G and 4A). Overall, the long-chain PFAS that contain six or more carbons are significantly more toxic than the short-chain PFAS that contain five or fewer carbons, which has been demonstrated in other studies (Figure 4B). Long-chain PFAS are excreted at a slower rate (therefore accumulate to a higher degree than short-chain PFAS) due to higher binding affinities to serum albumin and fatty acid-binding proteins, which may contribute to variation in protein-interaction kinetics and receptor activation between PFAS of varying chain lengths (Fenton et al. 2021; Jackson et al. 2021; Wolf et al. 2008; Zhang et al. 2013). This variation in toxicity is consistent with our results in *C. elegans*. Exposure to significantly higher doses of the three short-chain PFAS were required to elicit the same response (50% growth) compared to the long-chain perfluoroalkyl substances PFOSA, PFOS, PFNA, and PFOA. We observe the same result when we compare the per- and polyfluorether substances, GenX and PFEESA: GenX (C6) is significantly more toxic than PFEESA (C4) (Figure 4). However, it is critical to note that these two PFAS also differ by functional group, and that GenX is a branched molecule (Figure 1). Compellingly, we also observe that there is variation among strains when we compare susceptibility to long-vs short-chain PFAS, such as PFOA (C8) to PFBA (C4) and PFNA (C9) to PFPeA (C5) (Figure 6E, F). It is possible that genetic variation contributes to variation in uptake or elimination kinetics of PFAS in *C. elegans.* PFAS levels observed among humans varies widely even within populations (Fenton et al. 2021). One hypothesis is that variation in serum half-lives in humans is due to variation in transport proteins, particularly the organic anion transporters. Variation in transporter function, particularly transporters involved in renal clearance of PFAS, may result in particularly sensitive individuals. However, transport mechanisms are largely unknown as only nine renal uptake transporters have even been investigated for their role in PFAS transport in humans, and only in a limited number of PFAS (Niu et al. 2023). The *C. elegans* genome encodes 348 solute carriers, including four organic transporters homologous to human transporters, which provides a compelling system to address this significant gap in knowledge of the role in variation in PFAS uptake and elimination.

Exposure to a higher concentration of GenX is required to elicit the same adverse effect size as PFOA in *C. elegans*, which is consistent with some developmental studies in mouse and zebrafish (Blake et al. 2020; Gaballah et al. 2020; Satbhai et al. 2022). GenX has been described as the “regrettable replacement” of PFOA because an estimated 1.5 million people have been exposed to GenX since contamination of the largest watershed in the state of North Carolina, with detrimental health outcomes due to inadequate safety assessment (McCord and Strynar 2019; Pétré et al. 2022). Our results suggest that mechanisms of toxicity may vary, because the response among strains when we compare PFOA and GenX are significantly different (Figure 6G). This was expected, as GenX and PFOA greatly vary in their structural attributes, which may affect uptake and elimination kinetics, as well as variation in binding affinity to nuclear receptors. We measured equivalent bioconcentration factors between PFOA and GenX in *C. elegans*, which may suggest variation in modes of action contrary to variation in uptake or elimination kinetics, though there was variation among the GenX samples (Figure S4, Table S4). A few studies have demonstrated similar transcriptomic signatures of PFOA and GenX (Blake et al. 2022; Li et al. 2021). This uncertainty of specific molecular mechanisms of toxicity between these two PFAS, among many others, in *C. elegans* further supports future investigation of structure-specific mechanisms of PFAS toxicity.

In addition to variation in toxicity among PFAS, our results demonstrate that genetically divergent *C. elegans* strains vary in susceptibility to PFAS (Figure 5, Figure S3). Among all PFAS, there were roughly a 1.5 to 2.3-fold range in toxicity between the most sensitive and the most resistant strain (Figure 5, Table 1). This order of magnitude is lower than the traditional (though controversial) default intraspecies uncertainty factor of 10 that is currently used for assessing risk (Rusyn et al. 2022). However, the most sensitive adverse effect that we observed for each chemical varied by genotype. This suggests that toxicity testing in multiple genetic backgrounds, which is feasible in *C. elegans*, is important to take into consideration when assessing risk and is critical for future toxicity testing. For example, in only two of the ten PFAS (PFEESA and PFPeA) was the laboratory reference strain N2 the most sensitive strain. In fact, six different strains had the lowest EC50 values among the ten different chemicals tested. PFAS with the most variable response among strains were the perfluoroether acids GenX and PFEESA. Within these two PFAS as well as the short-chain PFCAs, PFPeA and PFBA, and the long-chain PFCA, PFNA, the strain DL238 was the most resistant to each exposure. DL238 is also resistant to starvation, which suggests that this strain harbors genetic variants that support stress resistance more generally (Webster et al. 2022). On the contrary, CB4856 is the most sensitive strain to respond to PFOS, PFNA, and PFOA, which suggests that this strain contains variants which contribute to sensitivity to exposure. The similarity of CB4856 in response to PFOA (C8), PFNA (C9), and PFOS (C8) supports a similar molecular mechanism of these long-chain PFAS. Among all strains, we saw no difference in *C. elegans* response to exposure when we compare the two long-chain PFCAs (PFOA and PFNA), nor between molecules that only vary by functional head group (PFOA and PFOS). This suggests that these long-chain PFAS may enact similar mechanisms of toxicity in *C. elegans*. We did however see variation among strains in their responses between PFNA and PFOS (Figure 6A), which are more similar in structure than PFOA and PFOS (barring functional group), which is supported by other studies that functional group contributes to PFAS mechanisms of action via variation in nuclear receptor activation (Yu et al. 2022; Zhao et al. 2023). Indeed, PFOS was significantly more potent than PFOA, which has also been observed in two recent *C. elegans* studies (Breton et al. 2023; Lin et al. 2022). This study was limited to twelve *C. elegans* strains, therefore screening a larger subset of genetically diverse wild *C. elegans* strains could illuminate variation among strains within and between PFOA and PFOS.

Genetic variation also contributes to variation in toxicity after exposure to short-chain PFAS, though this is driven by variation among strains in response to the sulfonamide, PFBSA (Figure 6B, C, and D). Interestingly, we observed no variation in toxicity among stains when we compared PFBA to PFPeA, PFBA to PFBS, and PFPeA to PFBS, which suggests that the sulfonamide functional group may have a particularly unique molecular mechanism of toxicity (Figure S5G, H, and K). Furthermore, there was no variation among strain response when we compared PFBSA to its long-chain complement, PFOSA (Figure S5P). This suggests that there is a particular mechanism of toxicity that is largely driven by this highly reactive functional group that is independent of the length of the molecule. Harnessing *C. elegans* natural variation has the potential to elucidate a sulfonamide-specific mechanism of toxicity. This is critical to investigate further because PFOSA and PFBSA are the most potent in this study and others, and FASAs are ubiquitous in the environment.

Overall, our results demonstrate that our culture system and imaging assay provide an effective approach to quantify the effects of PFAS exposure in *C. elegans*. We have demonstrated that not only is this approach useful to investigate toxicity across PFAS, but it can be used to investigate gene-by-environment interactions by harnessing the natural genetic variation among wild strains of *C. elegans*. Among the ten PFAS chemicals and twelve *C. elegans* strains investigated, we already determined that there is variation among strains in their response to specific PFAS structures. Our long-term goal is to harness the genetic tractability and incredible genomics toolkit of *C. elegans* to identify loci, genes, and variants that contribute to structure-specific mechanisms of PFAS toxicity. This is critical to understand, as it is impractical to quantitatively assess the safety of the >14,500 PFAS identified today. Currently, there are over 500 wild strains of *C. elegans* with distinct genome sequences and that come from unique geographic locations (Crombie et al. 2023). Therefore, screening additional wild strains of *C. elegans* from this unparalleled resource together with statistical and quantitative genetics and genome editing will allow for identification of specific genetic variants that contribute to variation in susceptibility to PFAS. This system can be used to identify mechanisms of toxicity due to specific structural attributes, systematically assessing and informing regulation of the thousands of other PFAS chemicals, and in doing so, improving human and environmental health.

## Acknowledgements

This research was funded by the National Institutes of Health (NIEHS, R01ES029930 to L.R.B. and P42 ES010356 to H.M.S.). T.C.L. was funded by a Charles. W. Hargitt Postdoctoral Fellowship (Duke Department of Biology) and F32-ES034954 (NIEHS). We would like to thank Dr. Erik Andersen (Johns Hopkins University) and his lab for providing the wild strains used in this study. We would also like to thank the members of the Andersen and Dr. Matt Rockman (New York University) laboratories for their helpful feedback throughout this study. We thank Drs. Linda Birnbaum, Justin Conley, and Joel Meyer for their constructive comments that significantly enhanced the manuscript.

