## Supplementary Materials for "Structure-specific variation in per- and polyfluoroalkyl substances toxicity among genetically diverse *Caenorhabditis elegans* strains"

Table S1. Test chemicals.

Table S2. Strains with significantly different EC50 values.

Figure S1. L1 lengths among twelve *C. elegans* strains**.**

Figure S2. Validate variation in growth among strains after 48 hr exposure to EC50.

Table S3. Strains with significantly different growth inhibition after 48 hr exposure to mean EC50.

Figure S3**.** Confirm estimated Effective Concentrations.

Figure S4. Bioconcentration factor of GenX and PFOA in *C. elegans.*

Table S4. Body burden and dosing solution concentrations of GenX and PFOA.

Figure S5. Variation in *C. elegans* susceptibility to structure-specific PFAS toxicity.

**Table S1**. Test chemicals used in this study.

| Chemical | Name | CASRN | MW (g/mol) | Vehicle | Concentrations Tested (µM) | Company |
|---|---|---|---|---|---|---|
| Perfluorooctane sulfonamide | PFOSA | 754-91-6 | 499.15 | MeOH | 0, 1.56, 3.125, 6.25, 12.5, 25, 50, 100 | SynQuest |
| Perfluorobutane sulfonamide | PFBSA | 30334-69-1 | 299.11 | MeOH | 0, 3.125, 6.25, 12.5, 25, 50, 100, 200 | SynQuest |
| Potassium perfluoro-1-octanesulfonate | PFOS | 2795-39-3 | 538.22 | MeOH | 0, 1.56, 3.125, 6.25, 12.5, 25, 50, 100 | Sigma |
| Perfluorononanoic Acid | PFNA | 375-95-1 | 464.08 | MeOH | 0, 3.125, 6.25, 12.5, 25, 50, 100, 200 | Sigma |
| Perfluorooctanoic Acid | PFOA | 335-67-1 | 414.07 | MeOH | 0, 15.6, 31.25, 62.5, 125, 250, 500, 1000 | Sigma |
| Ammonium perfluoro(2-methyl-3-oxahexanoate) | GenX | 62037-80-3 | 347.08 | DI water | 0, 390.625, 781.25, 1562.5, 3125, 6250, 12500, 25000 | SynQuest |
| Perfluoro(2-ethoxyethane)sulphonic acid | PFEESA | 113507-82-7 | 316.1 | MeOH | 0, 781.25, 1562.5, 3125, 6250, 12500, 25000, 500000 | SynQuest |
| Perfluorobutanesulfonic acid | PFBS | 375-73-5 | 300.1 | MeOH | 0, 195.25, 390.63, 781.3, 1562.5, 3125, 6250, 12500 | SynQuest |
| Perfluoropentanoic acid | PFPeA | 2706-90-3 | 264.05 | MeOH | 0, 781.25, 1562.5, 3125, 6250, 12500, 25000, 500000 | SynQuest |
| Perfluorobutanoic Acid | PFBA | 375-22-4 | 214.04 | MeOH | 0, 781.25, 1562.5, 3125, 6250, 12500, 25000, 500000 | SynQuest |
| Sodium 2-perfluoromethoxyl-2,2-difluoroacetate | PFMOAA | 21837-98-9 | 202.1 | DI water | 0, 390.625, 781.25, 1562.5, 3125, 6250, 12500, 25000 | SynQuest |
| 6:2 Fluorotelomer sulfonate | 6:2 FTS | 59587-39-2 | 445.198 | MeOH | 0, 15.6, 31.25, 62.5, 125, 250, 500, 1000 | SynQuest |
| (Perfluorohexyl)acetic Acid | 6:2 FCA | 53826-12-3 | 378.09 | MeOH | 0, 15.6, 31.25, 62.5, 125, 250, 500, 1000 | SynQuest |

**Table S2.** Strains with significantly different $log_{10}EC50$ values (companion to Fig. 5).

| Treatment | Strain 1 | Strain 2 | Adjusted p-value[1] | |
|---|---|---|---|---|
| PFBSA | N2 | MY23 | 2.52E-03 | ** |
| PFBSA | N2 | CX11314 | 4.71E-02 | * |
| PFBSA | JT11398 | MY23 | 7.81E-03 | ** |
| PFOS | CB4856 | MY23 | 1.72E-03 | ** |
| PFOS | EG4725 | MY23 | 2.34E-02 | * |
| PFOS | LKC34 | MY23 | 4.42E-02 | * |
| PFNA | N2 | CB4856 | 2.40E-06 | **** |
| PFNA | CB4856 | DL238 | 4.09E-07 | **** |
| PFNA | CB4856 | ED3017 | 4.74E-06 | **** |
| PFNA | CB4856 | JU258 | 7.70E-06 | **** |
| PFNA | CB4856 | JT11398 | 1.11E-05 | **** |
| PFNA | CB4856 | MY23 | 3.54E-05 | **** |
| PFNA | CB4856 | JU775 | 5.13E-05 | **** |
| PFNA | CB4856 | CX11314 | 9.67E-05 | **** |
| PFNA | CB4856 | EG4725 | 1.13E-04 | *** |
| PFNA | CB4856 | LKC34 | 1.86E-04 | *** |
| PFNA | CB4856 | MY16 | 2.44E-04 | *** |
| PFOA | N2 | CB4856 | 2.61E-02 | * |
| PFOA | CB4856 | ED3017 | 1.14E-03 | ** |
| PFOA | CB4856 | JU258 | 2.44E-03 | ** |
| PFOA | CB4856 | JT11398 | 2.72E-03 | ** |
| PFOA | CB4856 | MY23 | 4.38E-03 | ** |
| PFOA | CB4856 | JU775 | 8.28E-03 | ** |
| PFOA | CB4856 | DL238 | 1.44E-02 | * |
| PFOA | CB4856 | CX11314 | 1.47E-02 | * |
| PFOA | CB4856 | EG4725 | 1.96E-02 | * |
| PFOA | CB4856 | MY16 | 4.54E-02 | * |
| GenX | N2 | DL238 | 4.78E-04 | *** |
| GenX | CB4856 | EG4725 | 6.66E-04 | *** |
| GenX | CB4856 | JU258 | 1.06E-03 | ** |
| GenX | CB4856 | ED3017 | 8.51E-03 | ** |
| GenX | CX11314 | EG4725 | 6.53E-04 | *** |
| GenX | CX11314 | JU258 | 1.04E-03 | ** |
| GenX | CX11314 | ED3017 | 8.36E-03 | ** |
| GenX | DL238 | EG4725 | 1.55E-07 | **** |
| GenX | DL238 | JU258 | 2.46E-07 | **** |
| GenX | DL238 | ED3017 | 2.15E-06 | **** |
| GenX | DL238 | JT11398 | 1.51E-03 | ** |
| GenX | DL238 | MY16 | 3.10E-03 | ** |
| GenX | DL238 | LKC34 | 5.79E-03 | ** |
| GenX | DL238 | JU775 | 1.52E-02 | * |
| GenX | ED3017 | MY23 | 2.51E-03 | ** |
| GenX | EG4725 | MY23 | 1.82E-04 | *** |
| GenX | EG4725 | JU775 | 1.74E-02 | * |
| GenX | EG4725 | LKC34 | 4.31E-02 | * |
| GenX | JU258 | MY23 | 2.93E-04 | *** |
| GenX | JU258 | JU775 | 2.64E-02 | * |

Note: Significant interactions (Tukey's HSD) between strain $log_{10}EC50$ values within each PFAS chemical. [1] Indicates within-chemical adjusted p-value of Tukey's HSD test. * p<0.05, ** p<0.01, *** p<0.001, **** p<0.0001

| Treatment | Strain 1 | Strain 2 | Adjusted p-value[2] | |
|---|---|---|---|---|
| PFEESA | N2 | DL238 | 1.12E-10 | **** |
| PFEESA | N2 | MY23 | 3.30E-05 | **** |
| PFEESA | N2 | CX11314 | 8.65E-04 | *** |
| PFEESA | N2 | JT11398 | 6.67E-03 | ** |
| PFEESA | N2 | JU775 | 8.07E-03 | ** |
| PFEESA | CB4856 | DL238 | 4.17E-10 | **** |
| PFEESA | CB4856 | MY23 | 1.47E-04 | *** |
| PFEESA | CB4856 | CX11314 | 3.64E-03 | ** |
| PFEESA | CB4856 | JT11398 | 2.52E-02 | * |
| PFEESA | CB4856 | JU775 | 3.00E-02 | * |
| PFEESA | CX11314 | DL238 | 1.82E-04 | *** |
| PFEESA | CX11314 | EG4725 | 4.84E-02 | * |
| PFEESA | DL238 | EG4725 | 5.97E-09 | **** |
| PFEESA | DL238 | JU258 | 7.04E-08 | **** |
| PFEESA | DL238 | ED3017 | 1.35E-07 | **** |
| PFEESA | DL238 | LKC34 | 7.54E-07 | **** |
| PFEESA | DL238 | MY16 | 7.85E-07 | **** |
| PFEESA | DL238 | JU775 | 1.72E-05 | **** |
| PFEESA | DL238 | JT11398 | 2.12E-05 | **** |
| PFEESA | DL238 | MY23 | 4.43E-03 | ** |
| PFEESA | ED3017 | MY23 | 4.87E-02 | * |
| PFEESA | EG4725 | MY23 | 2.61E-03 | ** |
| PFEESA | JU258 | MY23 | 2.78E-02 | * |
| PFPeA | N2 | DL238 | 3.68E-06 | **** |
| PFPeA | N2 | MY23 | 7.59E-04 | *** |
| PFPeA | CB4856 | DL238 | 3.70E-04 | *** |
| PFPeA | CX11314 | DL238 | 2.29E-02 | * |
| PFPeA | DL238 | MY16 | 8.26E-05 | **** |
| PFPeA | DL238 | EG4725 | 1.19E-04 | *** |
| PFPeA | DL238 | ED3017 | 1.89E-04 | *** |
| PFPeA | DL238 | LKC34 | 5.30E-04 | *** |
| PFPeA | DL238 | JU258 | 6.34E-04 | *** |
| PFPeA | DL238 | JU775 | 1.10E-02 | * |
| PFPeA | DL238 | JT11398 | 3.44E-02 | * |
| PFPeA | ED3017 | MY23 | 2.89E-02 | * |
| PFPeA | EG4725 | MY23 | 1.93E-02 | * |
| PFPeA | MY16 | MY23 | 1.41E-02 | * |
| PFBA | N2 | DL238 | 5.15E-06 | **** |
| PFBA | CB4856 | DL238 | 1.62E-05 | **** |
| PFBA | CX11314 | DL238 | 8.90E-03 | ** |
| PFBA | CX11314 | JU258 | 2.86E-02 | * |
| PFBA | DL238 | JU258 | 1.51E-07 | **** |
| PFBA | DL238 | MY16 | 6.25E-06 | **** |
| PFBA | DL238 | EG4725 | 1.52E-05 | **** |
| PFBA | DL238 | ED3017 | 1.57E-05 | **** |
| PFBA | DL238 | LKC34 | 4.62E-05 | **** |
| PFBA | DL238 | JU775 | 6.37E-04 | *** |
| PFBA | DL238 | JT11398 | 2.71E-03 | ** |
| PFBA | DL238 | MY23 | 2.73E-03 | ** |

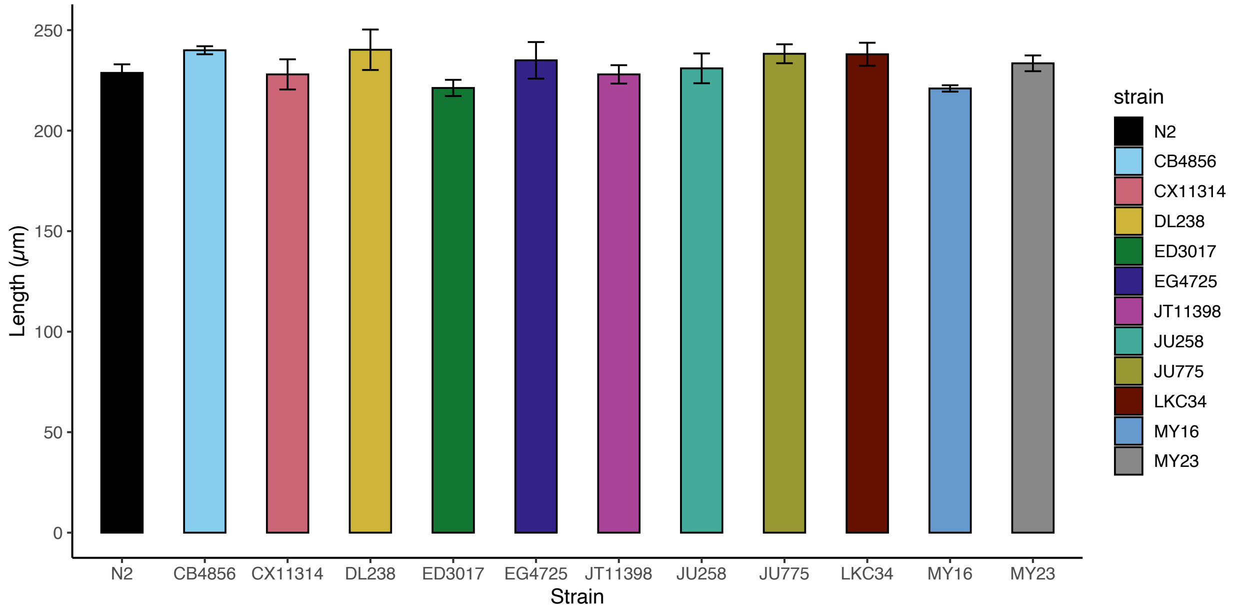


**Figure S1.** Lengths of twelve *C. elegans* strains during L1 arrest (prior to feeding or PFAS exposure)**.** L1 length was quantified 24 hr after bleaching and culturing in the absence of food. The mean length of L1 larvae in each well (an average of 22 $\pm$ 6 (standard deviation) worms per well) was determined ($n$ = 4 experimental replicates, p = 0.28, one-way ANOVA).

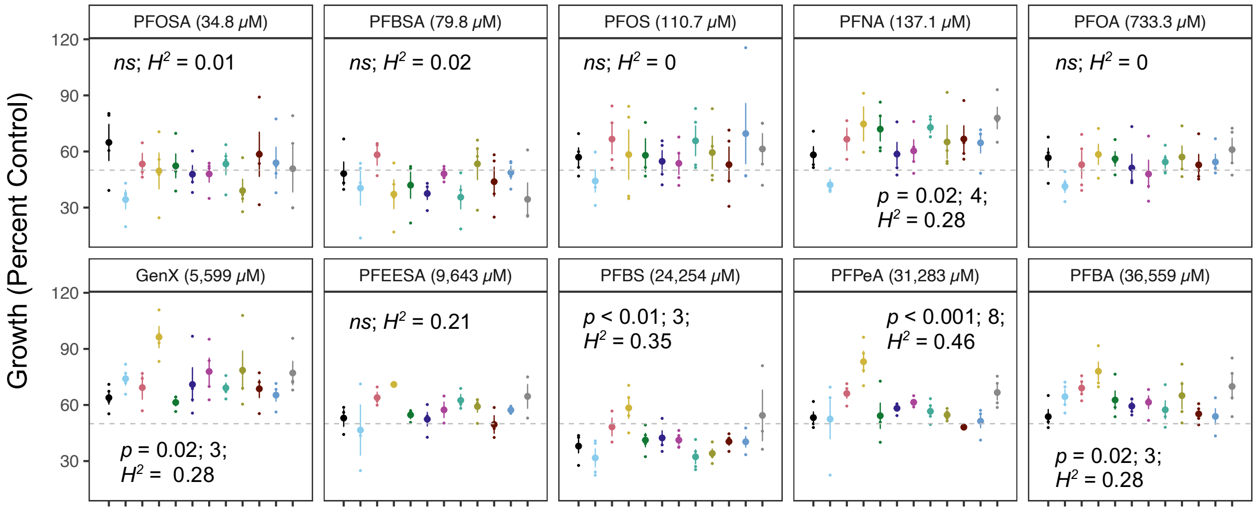


**Figure S2.** Validation of variation in growth among strains after 48 hr exposure to EC50. Each strain was exposed to each PFAS at the estimated EC50 (the concentration displayed in the title of each facet) and vehicle control for 48 hr, and mean length was determined. Strains are indicated by color. Percent control growth was calculated by dividing the mean length of the treated condition by mean control length for each strain within each chemical. The dashed line is at the expected relative length (50% control). Differences in EC values among strains were determined within each chemical (one-way ANOVA; *ns* = not significant). If there was significant variation detected, within-treatment differences in growth between each pair of strains were evaluated for significance using Tukey's HSD with correction for multiple comparisons. The number of significant pairwise interactions is displayed (significant pairwise interactions are listed in Table S3). Broad-sense heritability was estimated for each value (percent growth) as the trait ($H^2$) and is also displayed. Large dots are the mean growth per strain per treatment ($n$ = 3-4 experimental replicates per strain per treatment) and small dots reflect individual replicates. Error bars indicate standard error of the mean.

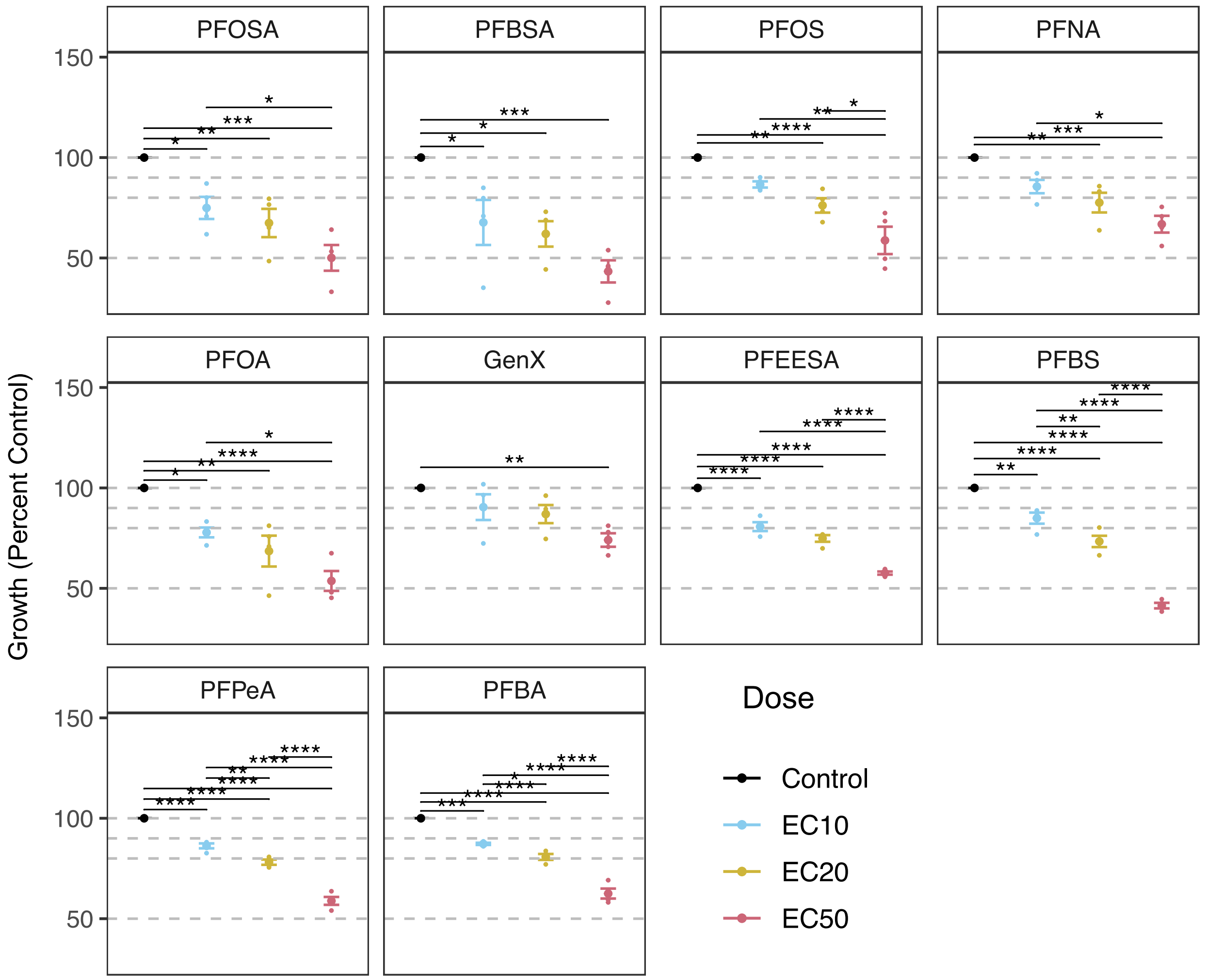


**Figure S3.** Validation of estimated Effective Concentrations. We exposed each of the twelve strains to the mean EC10, EC20, and EC50 concentration across all strains for each chemical, and we measured length at 48 hr to confirm 10, 20, and 50% growth inhibition at estimated EC values. Grey dashed lines indicate expected relative lengths based on EC values. We excluded wells that had less than five worms per well and calculated the mean length per treatment per replicate for each strain. An average of 20 individuals ± 7 (standard deviation) per well were analyzed. Each strain's mean length per replicate per treatment was then divided by each chemical's control value to obtain percent growth relative to control. The mean percent growth across all strains was calculated for each experimental replicate (*n* = 4 per chemical per concentraion). Error bars represent standard error of the mean. Variation in growth between each pair of exposure concentrations within a chemical was analyzed (two-way ANOVA, Tukey's HSD correction for multiple comparisons). Significant differences are indicated by asterisks (* $p < 0.05$, ** $p < 0.01$, *** $p < 0.001$, **** $p < 0.0001$).

**Table S3.** Strains with significantly different growth inhibition after 48 hr exposure to mean EC50 (companion to Figure S2).

| Treatment | Strain 1 | Strain 2 | Adjusted p-value[1] | |
|---|---|---|---|---|
| PFNA | CB4856 | DL238 | 4.35E-02 | * |
| PFNA | CB4856 | JU258 | 3.70E-02 | * |
| PFNA | CB4856 | MY23 | 7.92E-03 | ** |
| PFNA | CB4856 | ED3017 | 4.81E-02 | * |
| GenX | DL238 | N2 | 1.71E-02 | * |
| GenX | DL238 | ED3017 | 7.61E-03 | ** |
| GenX | DL238 | MY16 | 2.66E-02 | * |
| PFBS | CB4856 | DL238 | 9.98E-03 | ** |
| PFBS | DL238 | JU258 | 1.27E-02 | * |
| PFBS | DL238 | JU775 | 2.62E-02 | * |
| PFPeA | DL238 | N2 | 2.25E-03 | ** |
| PFPeA | DL238 | CB4856 | 1.56E-03 | ** |
| PFPeA | DL238 | ED3017 | 3.48E-03 | ** |
| PFPeA | DL238 | EG4725 | 1.93E-02 | * |
| PFPeA | DL238 | JU258 | 9.85E-03 | ** |
| PFPeA | DL238 | JU775 | 4.25E-03 | ** |
| PFPeA | DL238 | LKC34 | 2.17E-04 | *** |
| PFPeA | DL238 | MY16 | 9.55E-04 | *** |
| PFBA | DL238 | N2 | 2.63E-02 | * |
| PFBA | DL238 | LKC34 | 4.58E-02 | * |
| PFBA | DL238 | MY16 | 2.74E-02 | * |

Note: Significant interactions (Tukey's HSD) between strain growth (percent control) values within each PFAS chemical.

[1] Indicates adjusted p-value of Tukey's HSD test. * $p<0.05$, ** $p<0.01$, *** $p<0.001$

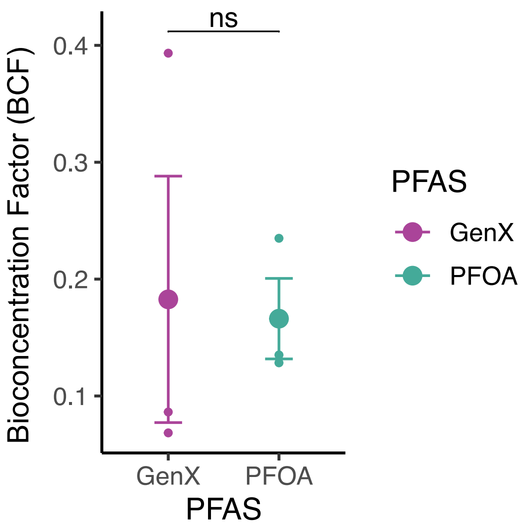


**Figure S4**. Bioconcentration factor of GenX and PFOA in *C. elegans*. LC/MS-MS was used to determine the bioconcentration factor (BCF) for GenX and PFOA, and BCF was calculated as [worm pellet]/[supernatant]. Worms accumulated GenX and PFOA and there was no difference in BCF between GenX and PFOA ($p = 0.91$, paired T-test, $n = 3$). Error bars represent standard error of the mean. See Table S4 for measured concentrations in worm pellet and supernatant.

**Table S4**. Mean (± S.E.) concentrations of GenX and PFOA were quantified via LC-MS/MS in the exposure medium (supernatant) and worm pellets to determine uptake of PFAS ($n$ = 3).

| | Treatment | GenX (µM) | PFOA (µM) |
|---|---|---|---|
| Supernatant | Control | 0.0 (± 0.0) | 0.9 (± 0.0) |
| | 500 µM GenX | 188.3 (± 96.2) | 0.9 (± 0.1) |
| | 500 µM PFOA | 0.1 (± 0.1) | 301.6 (± 46.7) |
| Worm Pellet | Control | 0.0 (± 0.0) | 0.0 (± 0.0) |
| | 500 µM GenX | 24.7 (± 9.9) | 0.1 (± 0.0) |
| | 500 µM PFOA | 0.0 (± 0.0) | 46.9 (± 1.31) |
| BCF | | 0.18 (± 0.11) | 0.17 (± 0.03) |

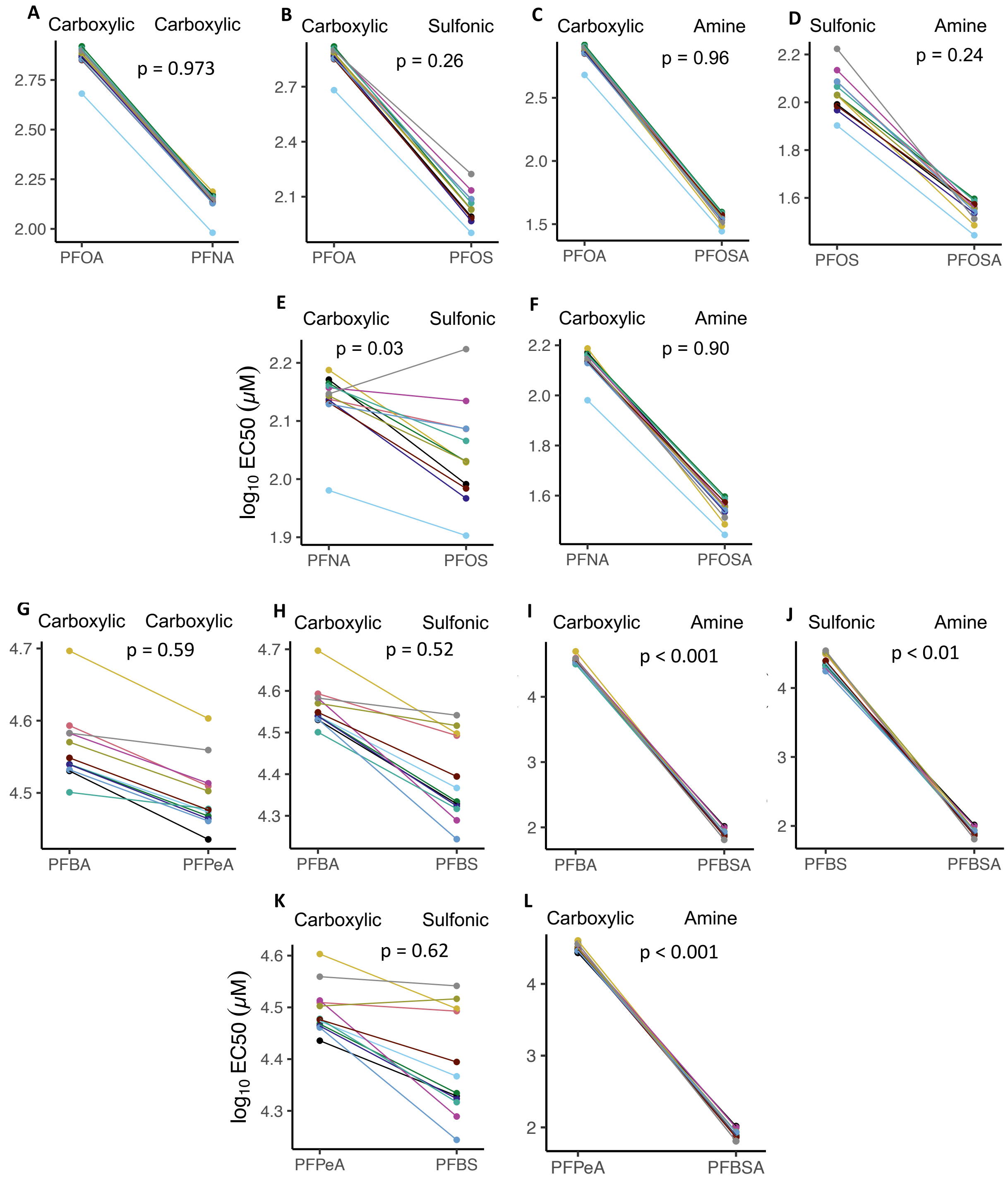
A
Carboxylic
Carboxylic
p = 0.973
PFOA
PFNA
B
Carboxylic
Sulfonic
p = 0.26
PFOA
PFOS
C
Carboxylic
Amine
p = 0.96
PFOA
PFOSA
D
Sulfonic
Amine
p = 0.24
PFOS
PFOSA
E
Carboxylic
Sulfonic
p = 0.03
PFNA
PFOS
F
Carboxylic
Amine
p = 0.90
PFNA
PFOSA
log10 EC50 (μM)
G
Carboxylic
Carboxylic
p = 0.59
PFBA
PFPeA
H
Carboxylic
Sulfonic
p = 0.52
PFBA
PFBS
I
Carboxylic
Amine
p < 0.001
PFBA
PFBSA
J
Sulfonic
Amine
p < 0.01
PFBS
PFBSA
K
Carboxylic
Sulfonic
p = 0.62
PFPeA
PFBS
L
Carboxylic
Amine
p < 0.001
PFPeA
PFBSA
log10 EC50 (μM)

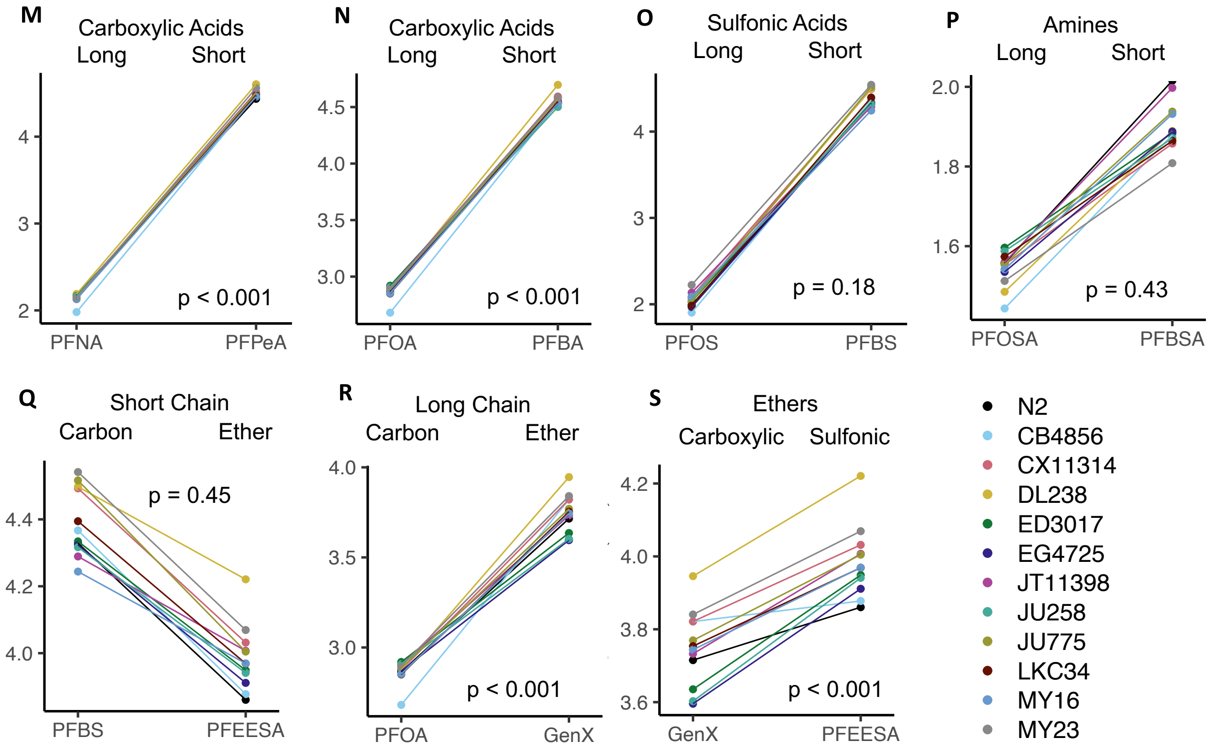


**Figure S5.** Variation in *C. elegans* susceptibility to specific PFAS structural attributes. Variation in toxicity ($log_{10}EC50$) among strains between structural classes of PFAS chemicals was analyzed (two-way ANOVA). Comparisons were made between long-chain PFAS that vary only by functional group: (A) the two carboxylic acids, (B, E) carboxylic vs sulfonic acids, (C, F) carboxylic acids vs amines, and (D) sulfonic acid vs amine. Comparisons were made between short-chain PFAS that vary only by functional group: (G) the two carboxylic acids, (H, K) carboxylic vs sulfonic acids, (I, L) carboxylic acids vs amines, and (J) sulfonic acid vs amine. Comparisons were made between long- and short-chain (M, N) carboxylic acids, (O) sulfonic acids, and (P) amines. Comparisons were made between carbon (non-ether) and ether for (Q) short-chain sulfonic acids (PFBS vs PFEESA) and (R) long-chain carboxylic acids (PFOA and GenX). (S) Comparison was made between two ethers (GenX and PFEESA) that vary in both length and functional group. Each strain is represented by the same color in each panel. Each dot represents the mean of each strain $log_{10}EC50$ (μM) (*n* = 4 experimental replicates per strain per treatment).
